# Metabolic plasticity and virulence of *Cryptococcus neoformans* are regulated by mitochondrial homeostasis

**DOI:** 10.64898/2026.02.06.704083

**Authors:** J. Alberto Patiño-Medina, Emma Camacho, Arturo Casadevall

## Abstract

The mitochondrion is a versatile organelle involved in diverse processes, such as cell death, metal homeostasis, plasma membrane and cell wall integrity, stress response, oxygen concentration, temperature, and metabolic adaptation, in addition to its role in generating energy. Consequently, mitochondrial fitness is essential for the pathogenicity of various organisms, including fungi. *Cryptococcus neoformans* is a fungal pathogen responsible for over 180,000 HIV-related deaths each year. In this study, we analyzed *C. neoformans* metabolic plasticity when grown with non-fermentable carbon sources and their impact on virulence and mitochondrial homeostasis. Growth on non-fermentable carbon sources increased thermotolerance, glucuronoxylomannan (GMX) content in the capsule, melanization rate, urease activity, biofilm formation, and virulence. Moreover, cells grown on non-fermentable carbon sources manifested increased mitochondrial number and activity. Conversely, mutants of the master regulator of mitochondrial biogenesis, the Hap complex, the catalytic subunit 1 of protein kinase A, or media supplementation with antioxidants, decreased the use of alternative carbon sources, capsule formation, melanin synthesis, urease activity, mitochondrial number, and resistance to both fluconazole and macrophage killing. Our results implicate mitochondrial homeostasis in virulence regulation via the PKA pathway, suggesting that targeting fungal mitochondrial homeostasis could be a therapeutic approach for cryptococcosis.

## Introduction

Metabolic plasticity is a key feature that enables microbial adaptation to different environments. For fungal pathogens, this feature is crucial for adapting to and surviving within the host, where nutrient availability includes alternative carbon sources beyond glucose and is, in part, dependent on mitochondrial function [1]. The mitochondrion is a versatile organelle, which, in addition to its role in energy production, is also involved in other fundamental processes such as metabolic adaptation, phenotypic changes, lipid metabolism and plasma membrane remodeling, metal and redox homeostasis, temperature and oxygen concentration adaptation, stress response, cell death, disease, and fungal pathogenesis, among others [1–3].

*Cryptococcus neoformans* is an important fungal pathogen that causes pulmonary and central nervous system infections in immunocompromised populations [4]. It is included on the World Health Organization’s Fungal Priority Pathogens List, released in October 2022. The mortality rate of cryptococcosis in patients with HIV infection remains alarmingly high, even with therapy, ranging from 41 to 61 %, while that of HIV-negative patients ranges from 8% to 20%, depending on the study [5]. Thus, it is essential to understand the molecular basis governing fungal physiology and, consequently, virulence potential.

*C. neoformans* is an obligate aerobic microbe [6], meaning that it relies on oxidative metabolism for energy to support its physiology, including primary metabolism and the expression of its virulence factors. Mitochondrial function is involved in fungal virulence [1], immune evasion [7], and antifungal resistance [8].

Additionally, the expression of fungal virulence factors, such as thermotolerance, capsule, melanization [9, 10], dimorphism [11], and biofilm formation [11] requires mitochondrial activity. Moreover, in relevant clinically fungal pathogens, the synthesis of some toxins also occurs specifically during oxidative metabolism [12–14].

Mitochondrial homeostasis is regulated by mitochondrial dynamics, which involves the biogenesis and fusion of mitochondria when energetic demand increases, as well as fission and mitophagy for mitochondrial clearance and recycling [15, 16].

Fungal mitochondrial biogenesis is governed by the transcriptional regulator Hap complex. This complex is evolutionarily conserved and is composed of a DNA-binding domain (Hap2, Hap3, and Hap5 subunits) that binds the CCAAT box, and an activation domain (Hap4/X subunit) [17]. The Hap complex transcriptionally regulates the expression of genes related to: 1) respiratory complexes and their assembly, 2) the TCA cycle and related metabolic pathways, 3) mitochondrial folding and import, 3) mitochondrial carriers and translocators, 4) mitochondrial fusion, fission, and mitophagy related genes [15, 16]. The Hap complex is active under aerobic/oxidative conditions, particularly during the use of non-fermentable carbon sources and glucose depletion. Conversely, it is repressed during fermentative conditions or when glucose is abundant [17].

The protein kinase A (PKA) pathway is a conserved signaling pathway in eukaryotic cells that regulates nutrient sensing, sugar and lipid metabolism, energy production, the cell cycle, and vesicular traffic, among other processes [18, 19]. In fungi, it also controls mating and virulence [20, 21]. Moreover, mitochondrial biogenesis and activity in fungi are also regulated by the PKA pathway [22–24].

This work analyzed the regulation of mitochondrial homeostasis and activity conferred by alternative carbon sources, as well as their impact on the expression of virulence factors in *C. neoformans* and on its virulence potential during host interaction.

## Results

### Non-fermentable carbon and inorganic nitrogen sources impact the growth rate and morphology *of C. neoformans*

In other fungi, fermentable carbon and organic nitrogen sources promote fermentative metabolism, whereas non-fermentable carbon and inorganic nitrogen sources promote oxidative metabolism [23–26]. To explore the effects of different nitrogen and carbon sources, we analyzed growth kinetics and morphology in response to these nutrients. We first investigated whether the concentration of the carbon source or the chemical nature of the nitrogen source impacted growth and morphology. The Yeast-Peptone-Dextrose (YPD) medium, which has an organic nitrogen source, provided optimal nutrients for growth, resulting in the highest cell density per milliliter at the end of the experiment, which was double that observed with Yeast Nitrogen Base (YNB) medium, which has an inorganic nitrogen source. YPD and YNB media both have a fermentable carbon source. Similar growth was observed between YNB and YNB low glucose media (20-fold reduction in glucose concentration compared with YNB); in contrast, YNB low-ammonium (10-fold reduction in ammonium concentration compared with YNB) and YNB low glucose and ammonium media resulted in reduced growth rate and cell density at the end of the experiment compared with the YNB condition (Fig. 1A). These results suggest that the chemical nature of the nitrogen source is more important than the glucose concentration in limiting growth. Subsequently, we analyzed glycerol, acetate, and ethanol as non-fermentable carbon sources. In general, the growth dynamics imply that *C. neoformans* needed a period to adapt to the new carbon source before achieving exponential growth; in addition, the final cell density achieved during stationary phase varied such that when ethanol was used as a carbon source was similar to that of the control YNB-glucose, followed by acetate, with a reduction of half of the cell density at the end of the experiment. Glycerol showed the slowest growth rate (Fig. 1B). Interestingly, we also observed changes in cell size and morphology. Cells from YNB lost their original spherical shape and had a rugose surface when compared with cells from YPD (Fig. 1C). Moreover, approximately 10 % of the population on YNB low-ammonium medium notably showed an increase in cell size (Fig. 1C). In contrast, the use of a non-fermentable carbon source resulted in decreased cell size when compared with YPD (Fig. 1C). These results suggest that *C. neoformans* can utilize both fermentable and non-fermentable carbon sources, as well as organic and inorganic nitrogen sources for growth and this metabolic plasticity impacts growth rate and morphology.

**Figure 1.**
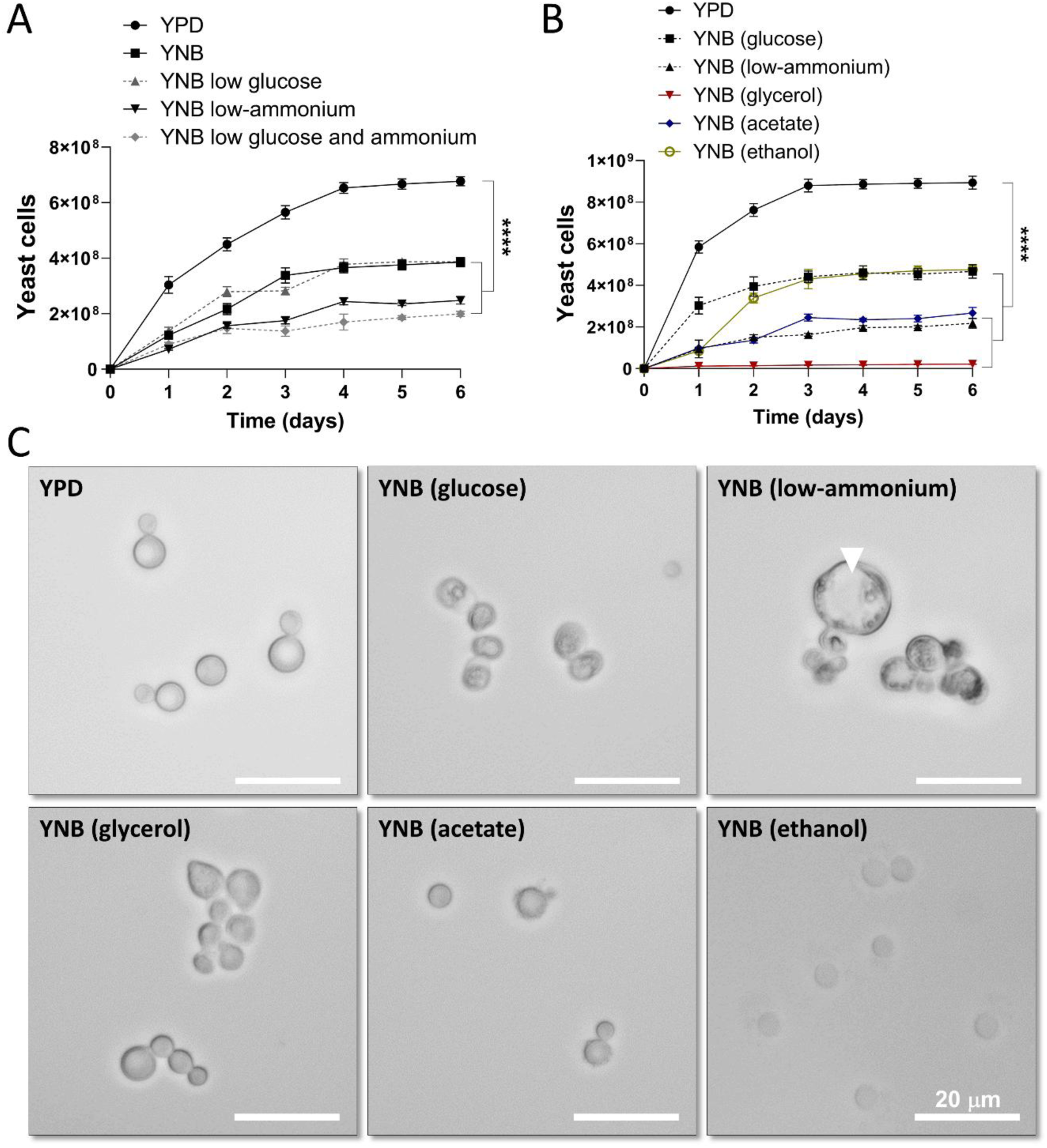
Effect of carbon and nitrogen source on the growth rate and cell morphology of *C. neoformans*. **(A)** Liquid growth kinetics of H99 strain in media with different concentrations of glucose or ammonium. **(B)** Liquid growth kinetics of H99 strain in media with fermentable or non-fermentable carbon, organic or inorganic nitrogen sources. Plots show the average of three biological replicates with technical duplicates, n=6, mean ± SD. One-way ANOVA with Tukey test, **** p<0.0001. **(C)** Representative microphotographs of the morphology of *C. neoformans* from panel B at day 6 with 100X magnification. The white arrowhead indicates an enlarged cell.

### The use of non-fermentable carbon and nitrogen sources regulates thermotolerance and capsule synthesis of *C. neoformans*

Non-fermentable carbon sources are metabolized in the mitochondria [27], and mitochondrial function is involved in fungal virulence [1]. Consequently, we evaluated the effect of carbon and nitrogen sources utilization on the expression of *C. neoformans* virulence factors.

Thermotolerance is a crucial fungal virulence factor for human infection, given the need for fungal cells to survive in mammalian hosts [28]. To analyze thermotolerance, yeast cells were grown in liquid media containing a non-fermentable carbon source and then inoculated into the same media (solid) at various incubation temperatures. Growth was similar between 30 and 37 °C; however, the use of non-fermentable carbon sources favored growth at 42 °C (Fig. 2A). These results suggest that oxidative metabolism contributes to adaptation and survival at high temperatures.

**Figure 2.**
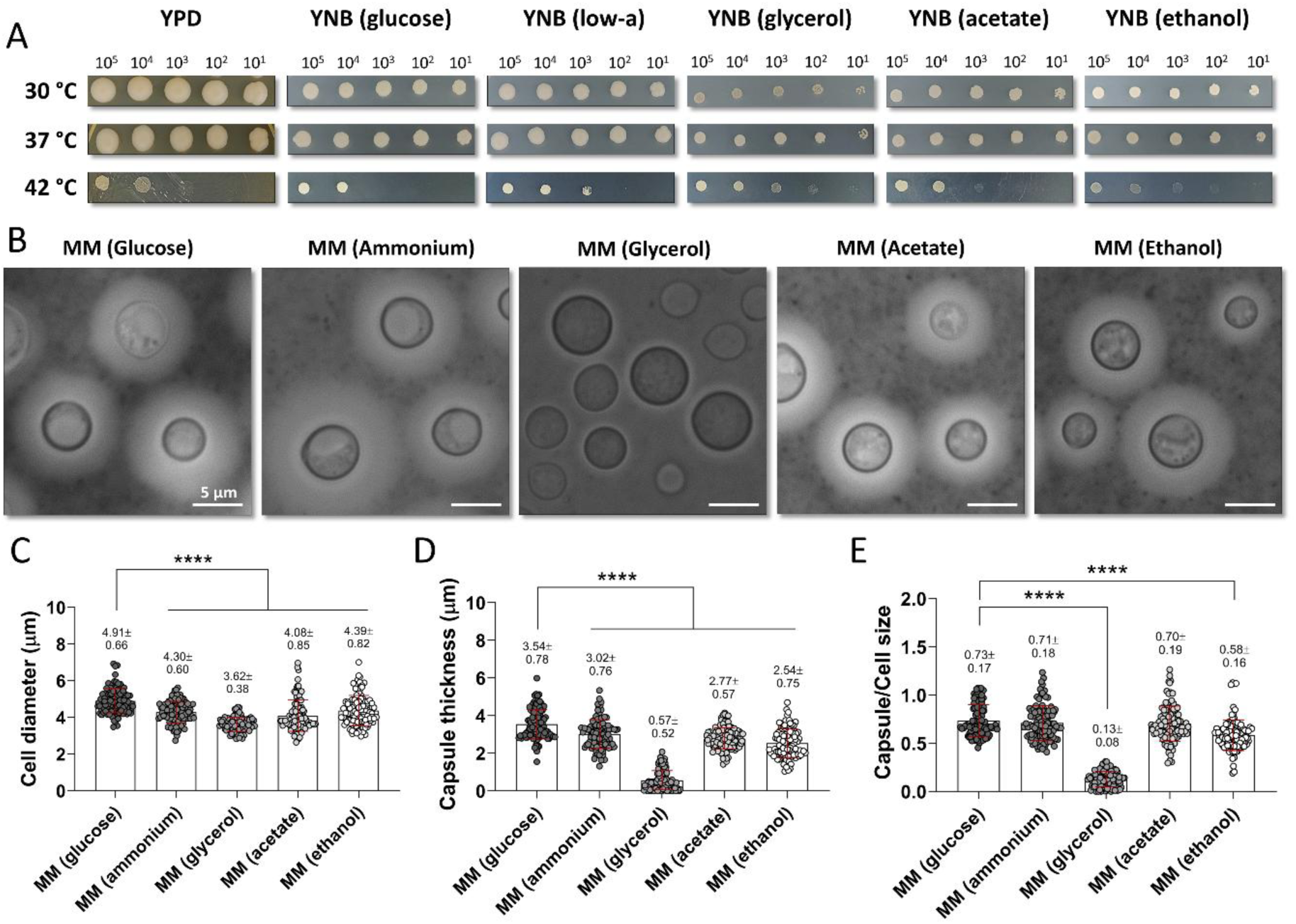
Using non-fermentable carbon sources supports capsule elaboration and increases thermotolerance of *C. neoformans*. **(A)** Representative photos at day 8 of H99 strain growth on solid media with non-fermentable carbon sources and different temperatures. **(B)** Representative microphotographs of India Ink staining of H99 strain grown on MM with non-fermentable carbon sources (day 4), 100X magnification. **(C)** Quantification of cell diameter (cell size) from panel B. **(D)** Quantification of the capsule thickness from panel B. **(E)** Quantification of the ratio between capsule and cell size from panel B. Plots show individual measurements of three biological replicates with technical duplicates, n=100, mean ± SD. One-way ANOVA with Tukey test, **** p<0.0001.

The capsule is essential for *C. neoformans* virulence [29]. To evaluate the influence of non-fermentable carbon sources (111 mM) on capsule production, yeast cells grown on the various media at 30 or 37°C were suspended in India Ink. Cell diameter (cell size) was quantified, showing that cells grown on non-fermentable carbon sources at 30 °C decreased their cell size by ∼30% (from ∼6 µm in YPD or YNB to ∼4 µm in glycerol, acetate, or ethanol). In contrast, in the YNB low-ammonium medium, ∼10% of the population exhibited cell enlargement (14.95 ± 0.63 µm), a 3-fold change compared with YPD (Fig. S1A and S1C). On the other hand, cells grown on YPD displayed cell size reduction at 37 °C when compared with 30 °C, while cells grown on non-fermentable carbon sources increased cell size at 37 °C compared with 30 °C (Fig. S1). At 37 °C, cell diameter measurements showed that cells grown in YNB-glucose and YNB low-ammonium exhibited a ∼20% increase in cell size compared to YPD. In contrast, cells from non-fermentable carbon sources showed a similar cell diameter to YPD (Fig. S1B and S1C). Quantification of capsule thickness at 30 °C showed a ∼50% reduction in the capsule of cells from YNB-glucose compared with YPD, as well as for cells from YNB low-ammonium and YNB-ethanol. Cells from glycerol or acetate media showed a slight reduction in capsule (Fig. S1A and S1D). Measurement of capsule thickness at 37 °C revealed a slight decrease in YNB-glucose, YNB low-ammonium, YNB-glycerol, and YNB-acetate media compared with YPD. Interestingly, cells grown on YNB-ethanol showed an increased capsule thickness at 37 °C compared with YPD. On the other hand, cells grown on YNB-glucose or YNB-ethanol exhibited an increase in capsule thickness at 37 °C compared with 30 °C (Fig. S1D). The ratio between capsule and cell size decreased in cells grown on YNB low-ammonium at both temperatures. In contrast, cells grown on YPD or YNB-ethanol increased at 37 °C compared with 30 °C (Fig. S1E). Together, these results imply that the chemical nature of nutrients and temperature influences cell size and capsule synthesis.

Since capsule synthesis is induced by nutrient starvation, yeast cells were grown in minimal media (MM) with an inorganic nitrogen source (13 mM) or with non-fermentable carbon sources (15 mM) at 30°C. Cell size measurements revealed a ∼20% reduction in all conditions compared with MM-glucose (Fig. 2B and 2C). Capsule thickness manifested a slight decrease when the MM contained an inorganic nitrogen source; on the other hand, when the MM contained acetate or ethanol, the capsule thickness was reduced by ∼30%. For MM-glycerol, the capsule thickness was dramatically reduced (Fig. 2B and 2D). The ratio between capsule and cell size of cells from MM-ammonium and MM-acetate was similar to that of cells grown in MM-glucose. However, in MM-glycerol and MM-ethanol, a reduction of ∼80% and ∼20% was observed, respectively (Fig. 2E). These results suggest that non-fermentable carbon sources can be used for capsule synthesis, which might influence capsule properties.

### The utilization of non-fermentable carbon sources alters cell-wall and capsule components of *C. neoformans*

To determine whether the carbon or nitrogen source influenced the cell wall and capsule, we quantified chitin and glucuronoxylomannan (GXM) content at 30 or 37 °C using fluorescence microscopy. The results at 30 °C showed that when *C. neoformans* used non-fermentable carbon sources, the GXM fluorescence intensity significantly increased 3.5 times compared to YPD (Fig. S2A and S2C); moreover, chitin fluorescence intensity was increased in YNB low-ammonium, YNB-acetate, and YNB-ethanol compared to YPD (Fig. S2A and S2D). At 37 °C, the GXM fluorescence intensity significantly increased in all conditions compared to 30 °C, whereas chitin fluorescence intensity significantly increased in YNB-glucose and YNB low-ammonium (Fig. S2B and S2C). Additionally, cells from YNB low-ammonium exhibited a significant GXM fluorescence intensity increase compared to YPD at 37 °C (Fig. S2B and S2C); moreover, chitin fluorescence intensity was significantly increased in YNB-glucose, YNB low-ammonium, and YNB-ethanol, while YNB-glycerol was decreased compared to YPD (Fig. S2B and S2D). These results imply that temperature, non-fermentable carbon sources, and inorganic nitrogen sources promote the levels of cell wall and capsule components.

### Non-fermentable carbon sources increased melanization rate, urease activity, and biofilm formation

Following the analysis of virulence factors. Yeast cells grown on different media were evaluated for melanization rate and urease activity. Cells grown on non-fermentable carbon sources showed a higher melanization rate at both temperatures, while higher urease activity at 37 °C compared to YPD (Fig. 3A and 3B). Interestingly, at 37 °C, melanization was delayed, and urease activity was downregulated at 3 h compared to 30 °C, but later reached similar levels. To analyze biofilm formation, cells from YPD were inoculated into the different YNB media. Cells in YNB, YNB low-ammonium, and YNB-glycerol exhibited an increase in biofilm formation at 30 °C. In contrast, cells in YNB-acetate and YNB-ethanol showed decreased biofilm formation at 37 °C (Fig. 3C). These results indicate that melanization, urease activity, and biofilm formation were enhanced when cells utilized non-fermentable carbon sources.

**Figure 3.**
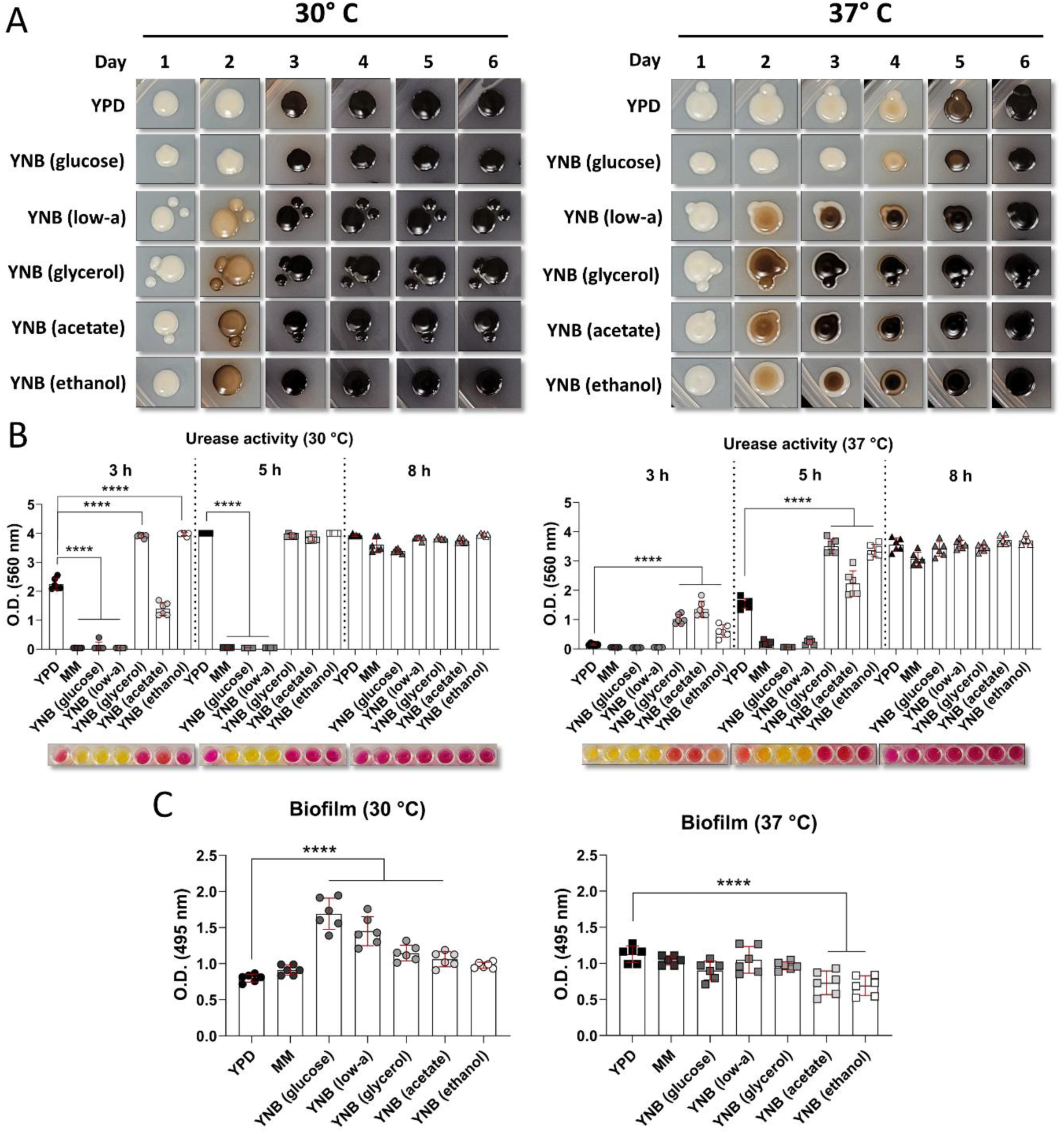
Using a non-fermentable carbon source increases the melanization rate, urease activity, and biofilm formation of *C. neoformans*. (**A**) Representative photos of melanization over time of H99 strain grown with non-fermentable carbon sources and spotted onto MM+L-DOPA. **(B)** Urease activity over time of H99 strain grown with non-fermentable carbon sources and inoculated into urea medium. **(B)** Biofilm formation of H99 strain in the presence of non-fermentable carbon sources. Plots show individual measurements of three biological replicates with technical duplicates, mean ± SD, n=6. One-way ANOVA with Tukey test, **** p<0.0001.

### Non-fermentable carbon sources increased mitochondrial biogenesis and activity of *C. neoformans*

Our results with non-fermentable carbon sources suggested that virulence factor expression is driven by increased mitochondrial function. To test this hypothesis, cells grown in the different carbon sources were stained with MitoTracker Red to assess mitochondrial number and activity. Mitochondrial localization within the cells was similar over time and across all conditions (Fig. 4A). However, the number of mitochondria per cell increased in cells grown with non-fermentable carbon sources compared to glucose, indicating enhanced mitochondrial biogenesis (Fig. 4B). Additionally, quantification of the fluorescence intensity revealed increased signal from cells grown with non-fermentable carbon sources, consistent with increased mitochondrial activity (Fig. 4C). Taken together, these results showed that non-fermentable carbon sources positively regulated mitochondrial biogenesis and activity.

**Figure 4.**
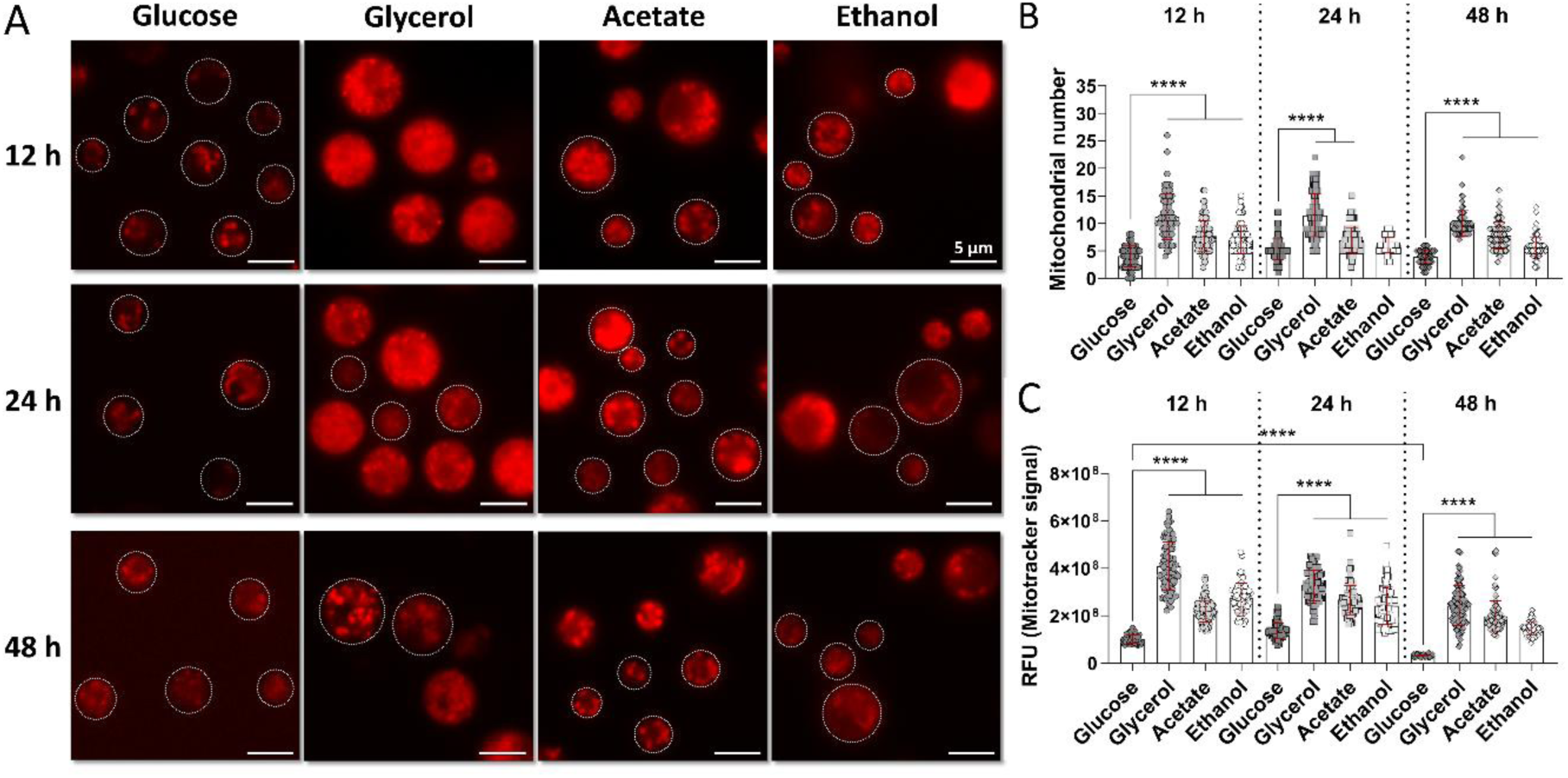
Using a non-fermentable carbon source by *C. neoformans* increases mitochondrial biogenesis and activity. **(A)** Representative microphotographs of H99 strain grown on non-fermentable carbon sources and stained with MitoTracker Red, 100X magnification. White circles indicate the shape of the yeast cell. **(B)** Quantification of mitochondria per cell over time. **(C)** Quantification of mitochondrial activity (fluorescent intensity) over time. Plots show individual measurements of three biological replicates with technical duplicates, n=100, mean ± SD. One-way ANOVA with Tukey test, **** p<0.0001.

### Mitochondrial biogenesis of *C. neoformans* is positively regulated during macrophage interaction

Metabolic plasticity is essential for fungal survival during host interactions. We therefore investigated how organelles involved in energy production, mitochondria and peroxisomes, were regulated in the context of pathogenesis. To test this scenario, mitochondria or peroxisomes [29] were stained in yeast cells before and after macrophage challenge. Interestingly, we observed that mitochondrial number increased significantly after macrophage interaction, whereas peroxisomal abundance remained unchanged (Figure 5A and 5B). The observations suggested that enhanced mitochondrial function promotes fungal survival during interaction with macrophages. To test this, macrophages were infected with yeast cells grown on non-fermentable carbon sources, and phagocytic index and fungal survival (CFUs) were assessed. Cells grown on non-fermentable carbon sources were phagocytosed less efficiently than cells grown on glucose. In contrast, cells grown in YNB were phagocytosed more efficiently than those grown on YPD (control) (Fig. 5C and 5D). Moreover, cells grown on non-fermentable carbon sources exhibited increased survival compared to YPD-grown cells (Fig. 5E). Therefore, to link the role of mitochondrial function with the ability of *C. neoformans* to use nutrients from tissues, we employed a condition that mimics host niches, agar made with liver or heart nutrients. We inoculated liver or heart tissue agar with H99 strain at 30 or 37 °C and stained mitochondria (Figure S3). The results showed that cells grown on tissue agar had mitochondrial numbers similar to those grown on MM-glucose at both temperatures (Figure 5F). However, cells grown on tissue agar exhibited increased mitochondrial function at both temperatures (Figure 5G). Together, these findings suggest that mitochondrial biogenesis and activity are positively regulated during host interaction, and that mitochondrial function is needed to metabolize nutrients from host niches.

**Figure 5.**
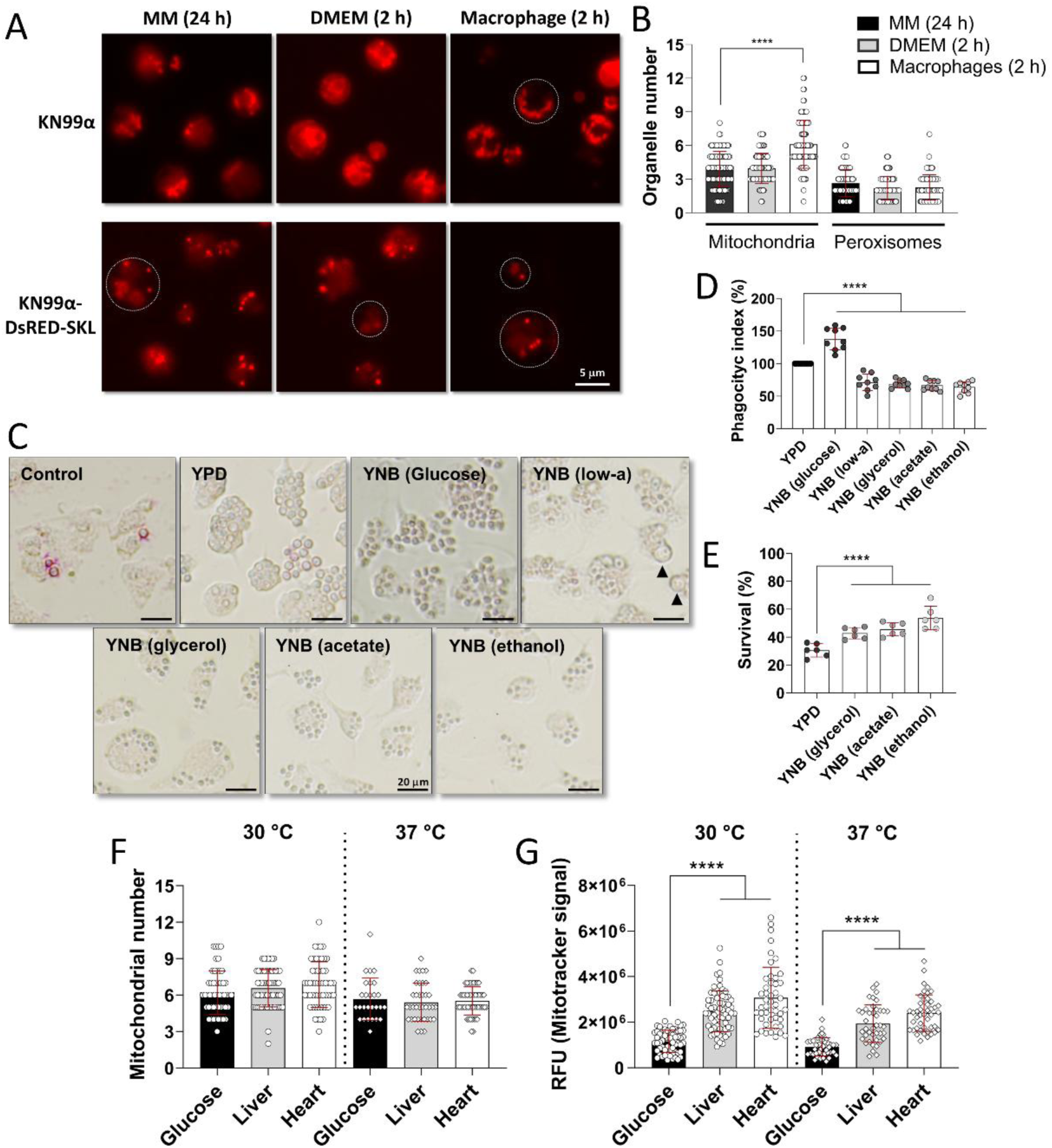
*C. neoformans* enhanced survival and resistance to phagocytosis through increasing mitochondrial biogenesis and activity. **(A)** Representative pictures of mitochondria (KN99α strain) or peroxisomes (KN99α-DsRED-SKL strain) stained from cells before and after interaction with bone marrow-derived macrophages (BMDMs), 100X magnification. White circles indicate the shape of the yeast cell. As a control, cells grown on MM, and cells without macrophages. **(B)** Quantification of mitochondria and peroxisomes per cell before and after interaction, n=50. **(C)** Representative microphotographs of phagocytosed yeast cells (H99 strain) grown on non-fermentable carbon sources, 100X magnification. Black arrowheads indicate enlarged cells. **(D)** Quantification of the phagocytic index from panel C, n=9. **(E)** Quantification of survival of yeast cells (H99 strain) grown on non-fermentable carbon sources after macrophage challenge, n=6. **(E)** Quantification of mitochondria per cell after growth on tissue-mimicking conditions, n=100. **(F)** Quantification of mitochondrial activity (fluorescent intensity) from panel E. Plots show individual measurements of three biological replicates with technical duplicates, mean ± SD. One-way ANOVA with Tukey test, **** p<0.0001.

### Mitochondrial biogenesis is essential for *C. neoformans* metabolic adaptation and virulence

The Hap complex governs mitochondrial biogenesis in fungi [17]. To analyze its role in *C. neoformans*, we evaluated mitochondrial homeostasis, metabolic adaptation, and virulence factor expression using a loss-of-function approach, employing the *hap3*Δ, *hap5*Δ, and *hapX*Δ strains, as well as their complemented counterparts [31].

When grown in YDP, MM, YNB-glucose, YNB-glycerol, YNB-acetate, or YNB-ethanol, the *hap* mutant strains showed growth defects compared to the WT strain (Fig. S4). On solid media, mutants exhibited impaired growth on non-fermentable carbon sources at 30°C, with more pronounced defects at 37°C, particularly in *hap3*Δ and *hap5*Δ (Fig. S5). The analysis of virulence-associated traits revealed that the *hap* mutant strains exhibited a thinner capsule (Fig. 6A and 6B), a reduced melanization rate (Fig. 6C), and reduced urease activity (Fig. 6D). In agreement with these findings, *hap*Δ mutant strains showed a dramatic reduction in mitochondrial number (Fig. 6E and 6F). Quantification of fluorescence intensity further showed a reduced mitochondrial activity in the mutants (Fig. S6). Growth of mutants in tissue-mimicking conditions showed delayed growth of *hap* mutants at 30 °C, while *hap3*Δ showed delayed growth at 37 °C (Fig S7). All defects were restored in the complemented strains. Taken together, these results demonstrate that the Hap complex directly regulates mitochondrial biogenesis and is essential for metabolic adaptation and for the expression of virulence factors in *C. neoformans*.

**Figure 6.**
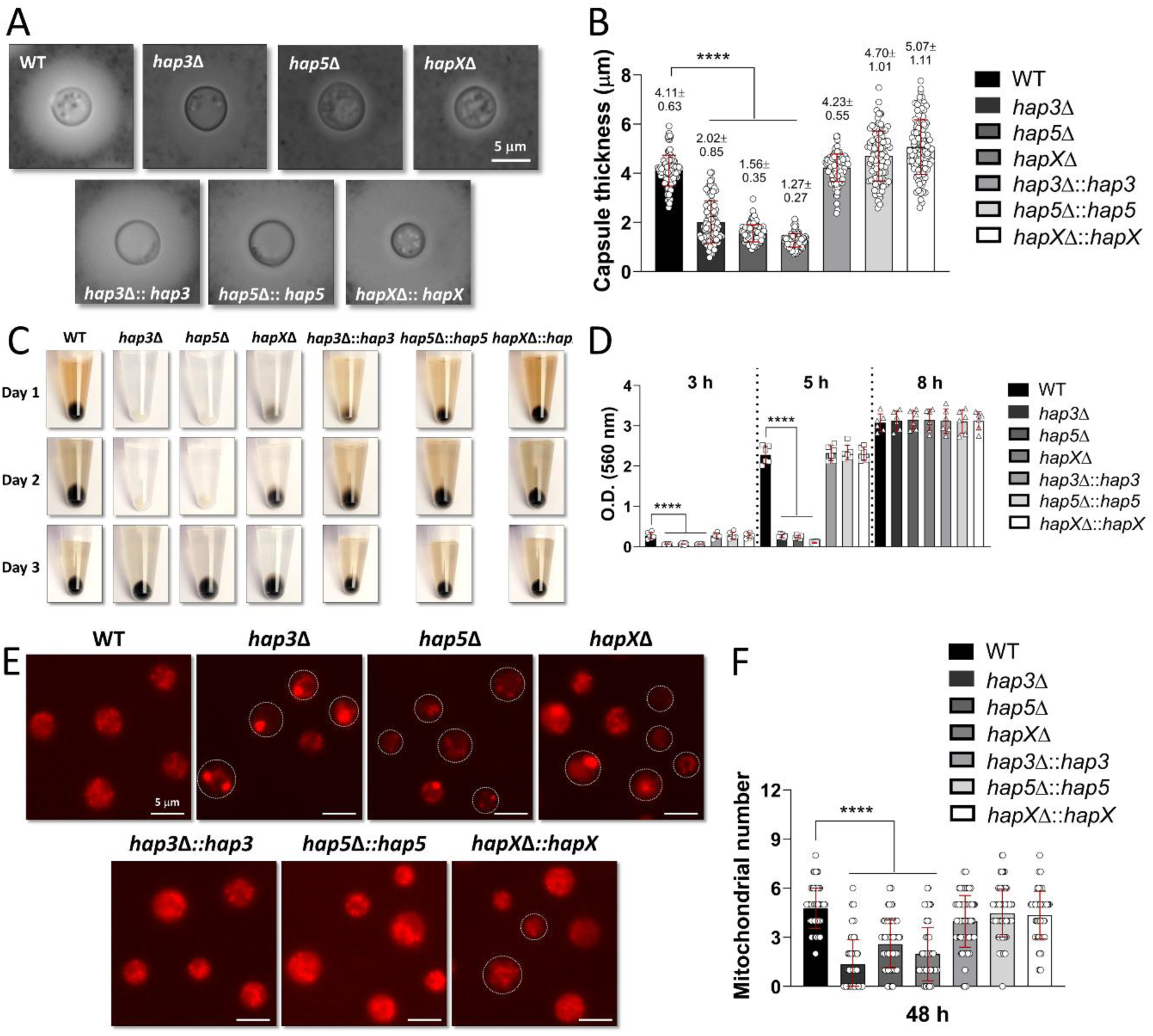
Mitochondrial biogenesis and activity are essential for the expression of virulence factors of *C. neoformans*. **(A)** Representative microphotographs of India Ink staining of the H99 (WT) and *hap*Δ mutant strains. **(B)** Quantification of the capsule thickness from panel A, n=100. **(C)** Representative photographs of melanization of the H99 and *hap*Δ mutant strains. **(D)** Urease activity over time of the H99 and *hap*Δ mutant strains, n=6. **(E)** Representative microphotographs at 48 h of the H99 and *hap*Δ mutant strains grown on MM and stained with MitoTracker Red, 100X magnification. White circles indicate the shape of the yeast cell. **(F)** Quantification of mitochondria per cell from panel E, n=100. Plots show individual measurements of three biological replicates with technical duplicates, mean ± SD. One-way ANOVA with Tukey test, **** p<0.0001.

### The PKA pathway contributes to mitochondrial biogenesis regulation in *C. neoformans*

In *S. cerevisiae*, mitochondrial biogenesis is regulated by the PKA pathway [22]. Hence, we investigated whether this regulatory mechanism is conserved in *C. neoformans*, where PKA activity is controlled by the Pka1 and Pka2 kinases [32]. Using two independent *pka1*Δ and *pka2*Δ mutant strains grown on MM, we analyze mitochondrial number and activity by fluorescence microscopy. The *pka1*Δ strain presented significantly fewer mitochondria than the WT over time, whereas the *pka2*Δ strain showed a comparable number of mitochondria to the WT (Fig. 7A and B). Consistently, the *pka1Δ* mutant strain showed reduced fluorescence intensity compared to WT, with the most pronounced reduction observed at the stationary phase (Fig. 7C). Additionally, *pka1*Δ mutants showed significant growth defects when grown on non-fermentable carbon sources (Fig. S8). Growth of mutants in tissue-mimicking conditions showed a delay in *pka1*Δ at 37 °C (Fig S7). These results demonstrate that the regulation of mitochondrial biogenesis and activity by Pka1 is conserved in *C. neoformans*.

**Figure 7.**
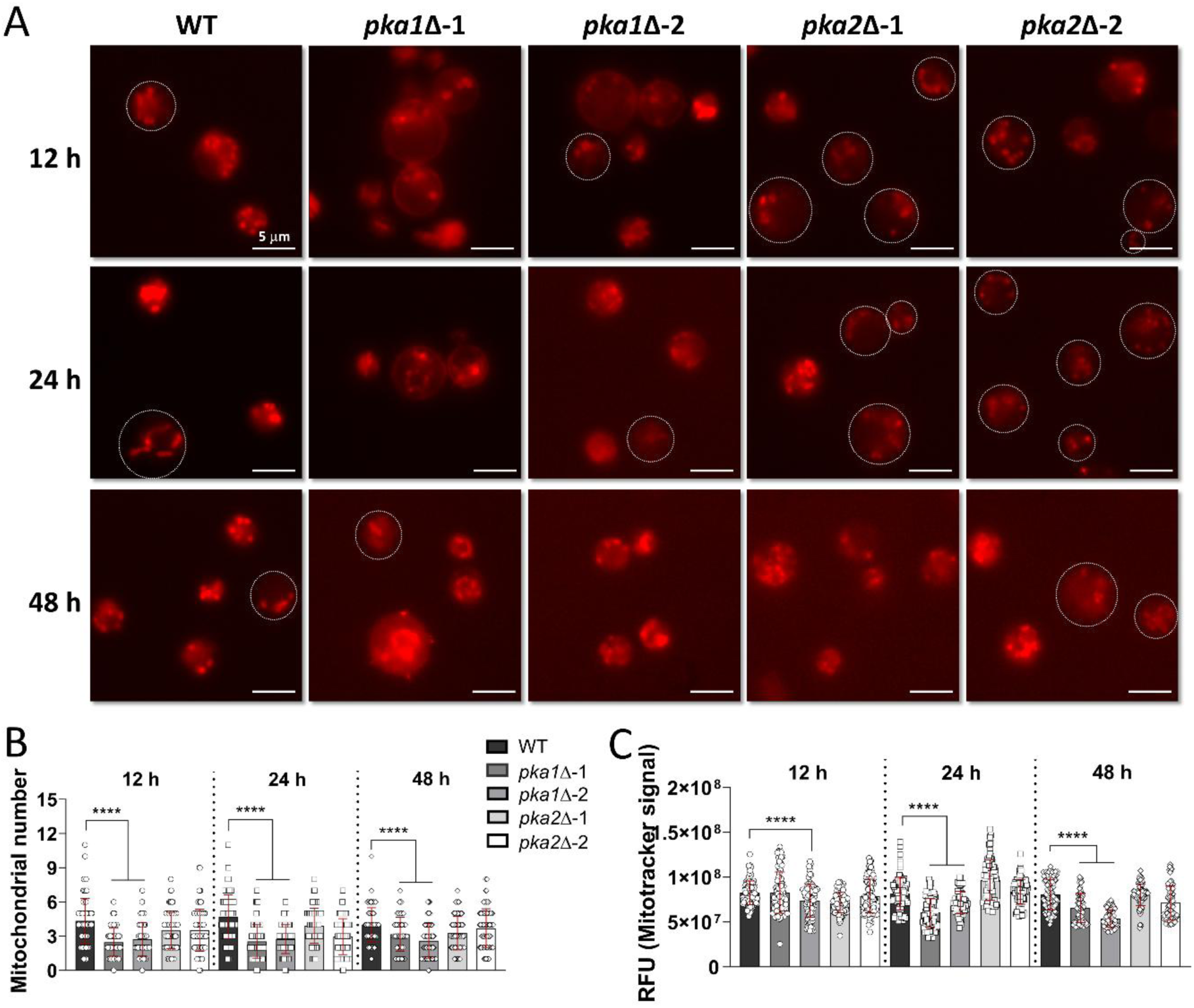
The PKA pathway positively regulates mitochondrial biogenesis and activity of *C. neoformans*. **(A)** Representative microphotographs over time of the H99 and *pka*Δ mutant strains grown on MM and stained with MitoTracker Red, 100X magnification. White circles indicate the shape of the yeast cell. **(B)** Quantification of mitochondria per cell from panel A, n=100. **(C)** Quantification of mitochondrial activity (fluorescent intensity) from panel A, n=100. Plots show individual measurements of three biological replicates with technical duplicates, mean ± SD. One-way ANOVA with Tukey test, **** p<0.0001.

### Hap5 and Pka1 are key for survival during macrophage interaction

Given that both the Hap complex and the PKA pathway regulate mitochondrial biogenesis and activity, as well as the expression of virulence factors in *C. neoformans*, we next investigated their contribution to host survival. Therefore, to demonstrate the genetic link between the Hap complex and the PKA pathway during the host response, we challenged *hap* and *pka* mutant strains with macrophages to assess fungal survival. These experiments revealed that Hap5 and Pka1 are key for fungal cell survival during a macrophage challenge (Figure S9). Together, these results demonstrate that the Hap complex contributes to *C. neoformans* virulence during macrophage challenge.

### Redox homeostasis is essential for virulence and antifungal susceptibility

Antioxidants inhibit melanization in *C. neoformans* [33]. We therefore hypothesized that redox imbalance affects mitochondrial fitness and the expression of other virulence factors. To test this, yeast cells grown on MM supplemented with glutathione or ascorbic acid were evaluated for capsule formation, melanization rate, and urease activity. Antioxidant treatment significantly reduced capsule thickness, melanization rate, and urease activity (Fig. 8A). ROS signaling plays a crucial role in maintaining mitochondrial homeostasis [34]. To investigate the role of ROS signaling in *C. neoformans* and its connection to mitochondrial biogenesis, we treated yeast cells with H_2_O_2_ and the antioxidants glutathione and ascorbic acid. At 24 h, antioxidant treatment reduced mitochondrial number, whereas H_2_O_2_ treatment had no detectable effect (Fig. 8B and 8C). In contrast, at 48 h, H_2_O_2_ treatment increased the number of mitochondria, while antioxidant-treated cells showed a similar number of mitochondria as the control (Fig. 8B and 8C). These results suggest that redox balance regulates mitochondrial biogenesis, with elevated ROS stimulating and reduced ROS repressing mitochondrial generation. Based on these results, we hypothesized that antioxidant treatment would affect antifungal susceptibility. To test this idea, yeast cells grown on MM supplemented with glutathione or ascorbic acid were treated with fluconazole or caspofungin.

**Figure 8.**
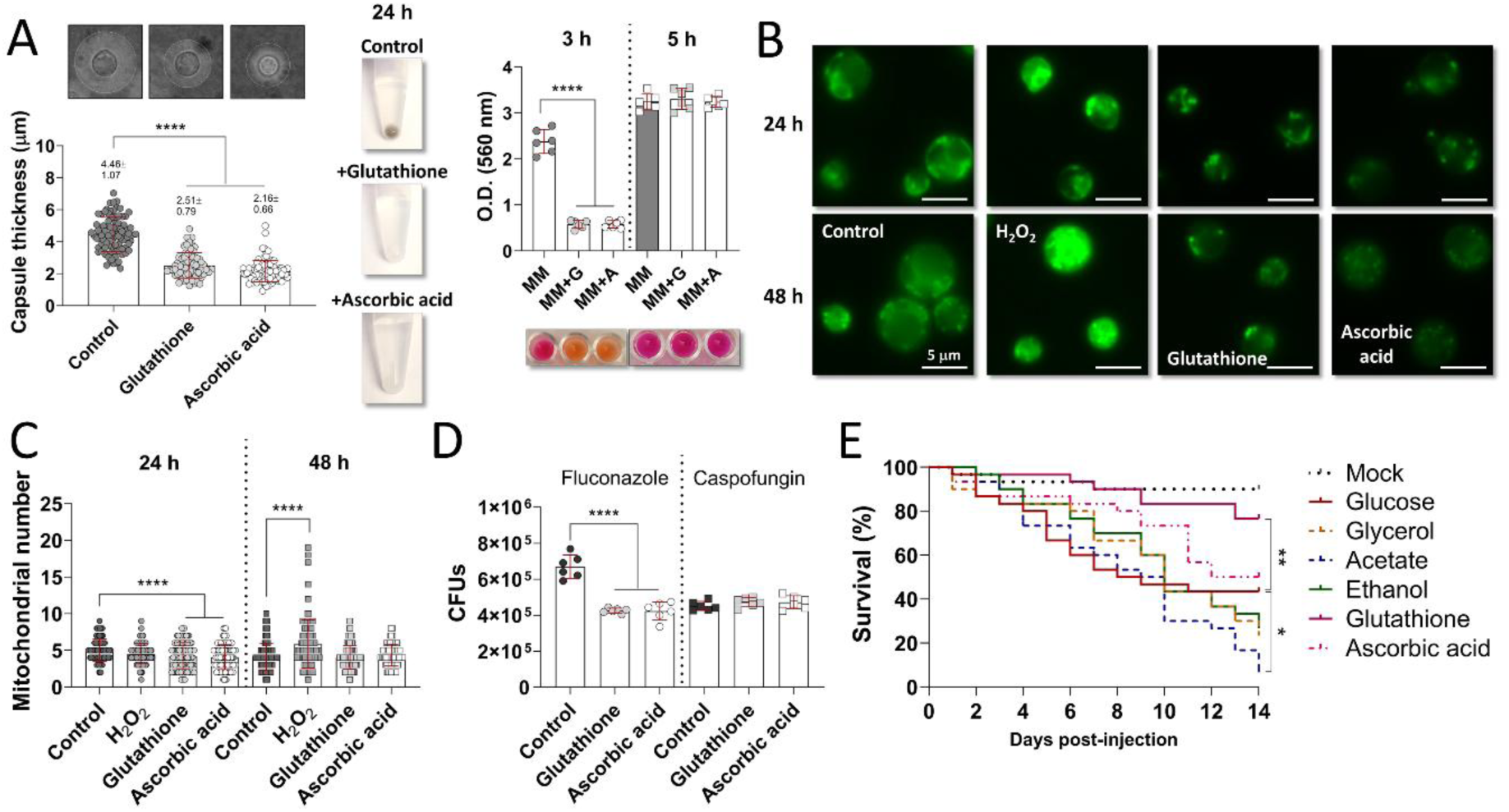
Redox homeostasis is key for mitochondrial biogenesis and virulence of *C. neoformans*. **(A)** Representative microphotographs of India Ink staining (left panel) of H99 strain treated with antioxidants, n=100. Representative photographs of melanization (center panel) of H99 strain treated with antioxidants. Urease activity (right panel) of H99 strain treated with antioxidants, n=6. **(B)** Representative microphotographs of MitoTracker Green staining of H99 strain treated with oxidant (H_2_O_2_) or antioxidants. **(C)** Quantification of mitochondria per cell from panel B, n=100. **(D)** Quantification of CFUs of H99 antioxidant-treated with antifungals, n=6. Plots show individual measurements of three biological replicates with technical duplicates, mean ± SD. One-way ANOVA with Tukey test, **** p<0.0001. **(E)** *Galleria* infection with H99 strain grown on non-fermentable carbon sources or treated with antioxidants, n=10. The plot shows the average of three independent experiments. Data was analyzed with the Mantel-Cox test, * p<0.05, ** p<0.01.

Antioxidant-treated cells were more susceptible to fluconazole, while susceptibility to caspofungin remained unchanged (Fig. 8D), consistent with the fact that echinocandins have no activity against *C. neoformans*. Finally, to test how virulence is modulated in an in vivo model, *Galleria mellonella* was injected with *C. neoformans* yeast cells grown on non-fermentable carbon sources or antioxidant-treated yeast cells. Results revealed that cells grown on non-fermentable carbon sources killed larvae at a higher rate than cells grown on glucose at the end of the experiment (Fig. 8E); however, only cells grown on acetate showed a statistically significant difference (p<0.0461) compared with those grown on glucose. In contrast, cells grown with antioxidants reduced death rate; however, only glutathione treatment (p<0.0021) abolished the virulence of *C. neoformans* (Fig. 8E). Together, these results suggest that ROS signaling is critical for maintaining mitochondrial fitness, fluconazole resistance, and virulence in *C. neoformans*.

### A model for mitochondrial role in *C. neoformans* virulence

Drawing on the findings of this study and those of other investigators cited in this paper, we synthesize a model for mitochondrial biogenesis and virulence regulation of *C. neoformans* (Figure 9).

**Figure 9.**
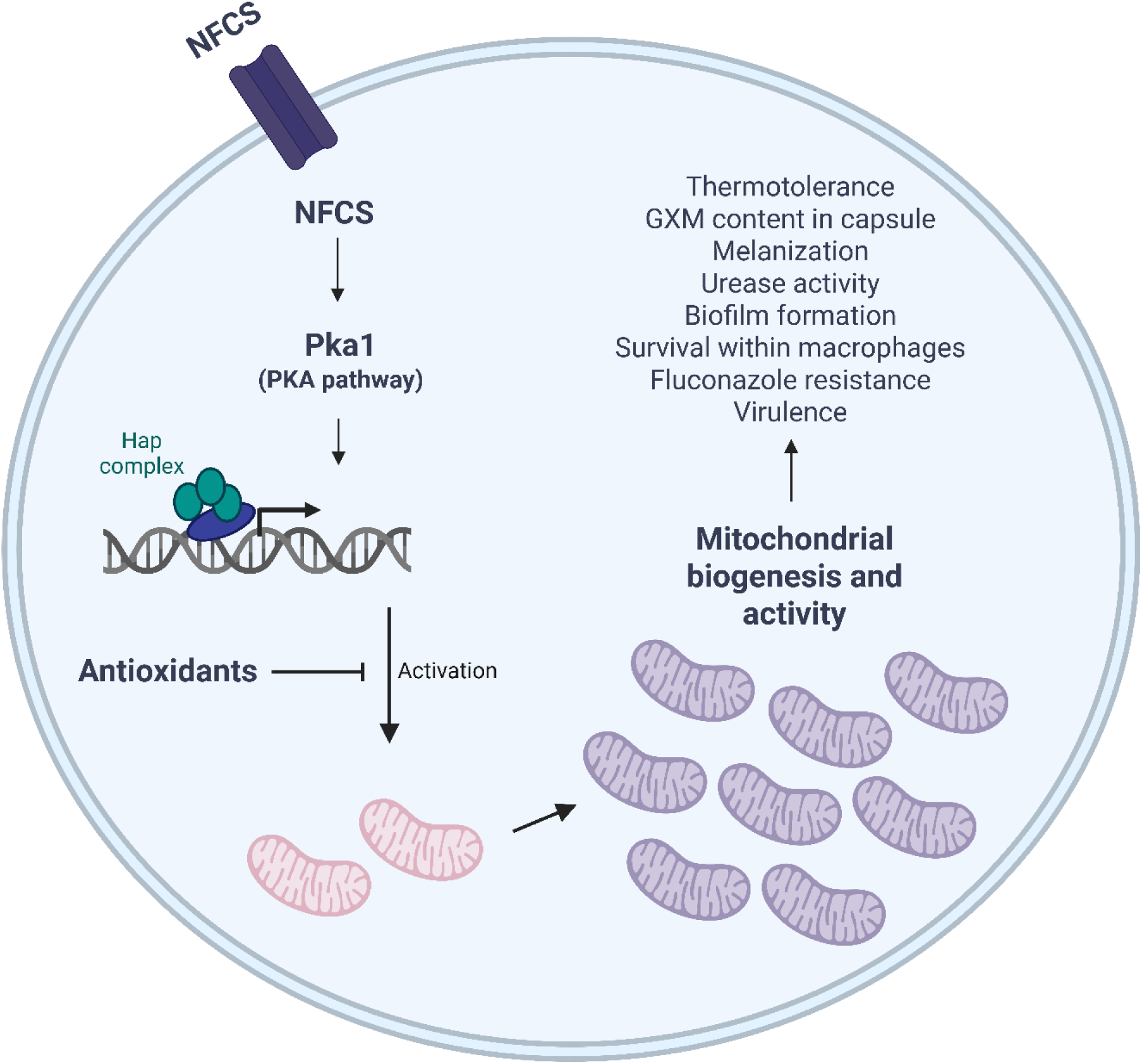
Model for mitochondrial biogenesis and virulence regulation in *C. neoformans*. Utilization of alternative carbon sources, including non-fermentable carbon sources (NFCS), increases mitochondrial number and enhances mitochondrial activity, a process mediated by the Hap complex through Pka1 (PKA pathway). Improved mitochondrial fitness results in upregulated expression of virulence factors, greater survival within macrophages, and increased fluconazole resistance. This regulatory network is disrupted by antioxidants, which interfere with ROS signaling. The figure was created using BioRender.

## Discussion

In this study, we examined how mitochondrial homeostasis regulates virulence in the human pathogen *C. neoformans*. Our results support the concept that mitochondria play key roles in cryptococcal metabolic plasticity, antifungal resistance, and virulence.

During infection, microbes must adapt to the host environment by using available nutrients, including carbon and nitrogen sources, and this adaptation involves mitochondrial function [35, 36]. Host-derived nutrients are a complex mix, including sugars, lipids, amino acids, nucleotides, metals, and vitamins. The competition for host defense (nutritional immunity) and microbial nutrient acquisition often determines the infection outcome [37, 38]. Although the specific nutrient types used by *C. neoformans* in tissue are unknown, they likely include tissue components such as degraded proteins, lipids, sugars, and metals. In this sense, glucose and iron metabolism are among the most studied processes in host-pathogen interactions [37]. Our data demonstrated that *C. neoformans* growth is supported by both fermentable and non-fermentable carbon sources and by organic and inorganic nitrogen sources in liquid and solid environments, whereas *C. neoformans* can grow using nutrients from liver and heart in tissue-mimicking conditions. These nutritional conditions led to differences in the growth rate and cell size of cryptococcal cells. While a higher growth rate contributes to invasion and dissemination, pathogen survival within the host is ultimately more critical for the establishment of infection [39]. Notably, a low concentration of ammonium sulfate, an inorganic nitrogen source, promotes the growth of larger cells under liquid aerobic conditions. Enlarged cell size leads to a titan cell phenotype, a mechanism for evading macrophage phagocytosis [40]. In contrast, the use of non-fermentable carbon sources significantly reduced cell size, with daughter cells exhibiting a 3.5 ± 0.30 µm cell diameter under these conditions. It has been proposed that smaller cells disseminate more effectively within the host [41].

The repertoire of virulence factors determines a microbe’s success during infection, and these usually function to promote survival in vivo, often by enabling immune evasion [39]. *C. neoformans* displays various such factors, including the capsule, melanization, urease activity, and biofilm formation. The capsule is primarily composed of the polysaccharide GMX and possesses immunomodulatory properties that help to evade host immune response [29]. Melanin confers resistance to drugs, oxidative stress, macrophage phagocytosis, and promotes brain dissemination [42, 43]. Urease activity facilitates survival in the acidic pH of phagolysosomes and supports brain dissemination [44]. Biofilm formation increases resistance to host immune mechanisms and antimicrobial therapy [45].

Our data show that growth in non-fermentable carbon sources increases tolerance to high temperatures, enhances GXM content in the capsule, augments melanization rate and urease activity, and promotes biofilm formation. The increased expression of these virulence factors is consistent with the low phagocytic index and higher survival observed when cells grown on non-fermentable carbon sources were challenged with macrophages, particularly due to the higher GXM content in the capsule and urease activity. In addition, cells grown on non-fermentable carbon sources were more virulent than those grown on glucose in the *G. mellonella* model. Interestingly, we also observed that chitin and GXM content were increased at 37 °C, suggesting that capsule and cell wall are remodeled at host temperature and in the presence of non-fermentable carbon sources. Future analyses are needed to elucidate this hypothesis. Moreover, thermotolerance conferred by non-fermentable carbon sources may be especially relevant at environmental temperatures above those of the host, as *C. neoformans* exhibited improved thermotolerance at 42 °C. Capsule synthesis in *C. neoformans* was repressed by glycerol, which also caused the lowest growth rate under the conditions tested in this study. Glycerol may exert a stronger control of the carbon catabolite repression (CCR) pathway than the other non-fermentable carbon sources analyzed, although further work is needed to test this hypothesis. In *C. neoformans*, Mig1, the transcription factor responsible for regulating CCR [38], controls the expression of amino acid, heme, and carboxylic acid biosynthesis, as well as genes associated with the TCA cycle, and the electron transport chain, all of which are involved in mitochondrial function. Moreover, mitochondrial function is co-regulated by Mig1/HapX under iron starvation, forming a regulatory network that is essential for *C. neoformans* virulence [46].

Non-fermentable carbon sources are metabolized in the mitochondria and positively regulate mitochondrial function. Consistent with this, we observed that the use of non-fermentable carbon sources increases mitochondrial number and activity in *C. neoformans*, thereby improving cellular metabolic fitness.

Mitochondria and peroxisomes are key organelles involved in energy production and in carbon and nitrogen metabolism [47]. Peroxisomal β-oxidation has also been reported to contribute to *C. neoformans* virulence [48, 49]. During macrophage interaction, however, our data revealed an increase in mitochondrial number but not in peroxisome number, suggesting that mitochondrial activation represents an early response to host interaction. In *C. albicans*, respiratory activity is required for filamentation within the phagosome, for escape from it, and for inducing phagocyte cell death [6]. Moreover, when *C. neoformans* was grown under tissue-mimicking conditions, mitochondrial function was enhanced.

Mitochondrial biogenesis is a central regulatory step in mitochondrial dynamics, and it’s critical for energy production, metal homeostasis (iron, copper, calcium, etc.), and mitochondrial inheritance. The Hap complex is a master regulator of mitochondrial biogenesis in fungi, one of its canonical functions [17, 50]. Our data revealed that deletion of Hap complex subunits in *C. neoformans* leads to a dramatic reduction in mitochondrial number and activity under nutrient-poor conditions, indicating that the Hap complex directly controls mitochondrial biogenesis. In *hap*Δ mutans that form the DNA-binding complex, we also observed defects in growth on non-fermentable carbon sources or liver/heart nutrients, thermotolerance, melanization rate, and urease activity. Furthermore, macrophage challenge revealed increased cell death in the *hap5*Δ mutant. Given that the Hap complex regulates iron homeostasis, capsule synthesis, and virulence in *C. neoformans* [31], our results are consistent with previous findings and provide new insights into the link between mitochondrial biogenesis, use of non-fermentable carbon sources, and regulation of other virulence factors (urease activity and biofilm formation).

The PKA pathway is also implicated in the regulation of fungal virulence, including *C. neoformans* [51]. The *C. neoformans* genome encodes two catalytic subunits of PKA, *pak1* and *pka2* [32]. Pka1 is required for mating, capsule formation, and melanin production. When investigating the role of the PKA pathway in mitochondrial homeostasis, we found that loss of *pka1* leads to reduced mitochondrial number and activity under nutrient-poor conditions, as well as growth defects on non-fermentable carbon sources or liver/heart nutrients, phenotypes similar to those of *hap*Δ mutants. In agreement with our findings, the PKA pathway regulates mitochondrial biogenesis and activity in other fungi such as *S. cerevisiae* [22] and *M. lusitanicus* [24]. In addition, macrophage challenge assays revealed increased cell death in *pka1*Δ mutant strains. Together, these results indicate that mitochondrial fitness and PKA signaling are essential for fungal metabolic adaptation and virulence. In *C. albicans*, mitochondrial metabolism of amino acids arginine, ornithine, and proline induces yeast-to-hyphal transition, the virulent morphology of *C. albicans*. This trait requires the activation of the cAMP-PKA pathway [35]. Similarly, in *Mucor lusitanicus*, cAMP-PKA signaling pathway is needed for mitochondrial function and virulence [24]. Additionally, the synthesis of fungal toxins requires oxidative metabolism and mitochondrial function, for example, candidalysin in *C. albicans* [11], mucoricin in *Rhizopus delemar* [12], or gliotoxin in *A. fumigatus* [13].

One subproduct of mitochondrial function is the generation of ROS. Excessive ROS can disrupt redox homeostasis and damage cellular components. However, ROS signaling plays an important role in fungal biology [52]. Our data show that antioxidant supplementation attenuates capsule synthesis, urease activity, mitochondrial biogenesis, fluconazole resistance, and virulence in *G. mellonella*, revealing that redox balance is essential for mitochondrial homeostasis and fungal virulence. Consistent with these findings, antioxidants inhibit melanization [33] and titan cell formation [53]. In *M. lusitanicus*, antioxidant supplementation has similarly been reported to attenuate mitochondrial function and virulence [24]. In contrast, ROS stimulation increased the secretion of the siderophore rhizoferrin, which promotes iron uptake and virulence [23]. In outbreak strains of *C. gattii*, ROS produced by macrophages promote the intracellular survival of a subpopulation of yeast cells, a phenotype conferred by mitochondrial tubularization (mitochondrial fusion) [54]. In addition, in *C. neoformans*, a low glucose concentration (0.05 %) enhanced mitochondrial function, thereby contributing to fluconazole resistance by increasing efflux pump activity [55]. Consistent with this observation, our data showed that antioxidant supplementation decreased fluconazole resistance, the opposite phenotype. The finding that the cell’s redox state affects fluconazole susceptibility suggests that in vivo and in vitro susceptibilities can differ and raises the intriguing possibility that antioxidant supplementation may enhance drug efficacy in patients. In that regard, vitamin C enhances the activity of antituberculosis drugs against *M. tuberculosis* [56]. Given that antioxidant supplementation decreased both virulence factor expression and fluconazole resistance, combining antioxidant treatment with existing therapeutic strategies may represent a synergistic approach for preventing and treating fungal infections.

In summary, mitochondrial homeostasis governs metabolic plasticity and virulence in *C. neoformans,* with the Hap complex regulating mitochondrial biogenesis through the PKA pathway. The mitochondrial-related effects reported here are protean, ranging from changes in thermal tolerance to alterations in virulence factor expression to differences in antifungal susceptibility, suggesting that manipulation of fungal mitochondrial expression is a fertile area for investigating new options to improve therapeutic outcomes.

## Materials and methods

### Strains and Culture Media

The different strains used in this study are listed in **Table S1**. Kronstad Lab kindly provided the *hap* mutant strains. Heitman Lab kindly provided the *pka* mutant and AI100 strains. For growth analysis, various media were used. YPD broth (Difco, NJ, USA), rich medium, with glucose as a fermentable carbon source and peptone as an organic nitrogen source, and minimal medium Yeast Nitrogen Base without amino acids and ammonium sulfate (YNB) (Difco). Ammonium sulfate (Sigma, Missouri, USA) was added as an inorganic nitrogen source, and glycerol (Sigma), acetate (Sigma), or ethanol (Sigma) as non-fermentable carbon sources, respectively.

Each liter of YPD medium contained 50 g of YPD broth (2% of glucose), while YNB contained 1.7 g of YNB base, 20 g (2%/111 mM) of glucose, and 40 mM of ammonium sulfate. For YNB low glucose concentration medium (YNB low glucose), glucose was at 0.1 %; for YNB low ammonium concentration medium (YNB low-Ammonium), ammonium sulfate was at 4 mM; for YNB low glucose and low ammonium concentration medium (YNB low glucose and ammonium), glucose was at 0.1 %, and ammonium sulfate was at 4 mM. For YNB-glycerol, YNB-acetate, or YNB-ethanol media, each non-fermentable carbon source was added at 111 mM instead of glucose. Nitrogen and carbon sources were used at equimolar concentrations to the standard YNB. When solid media were required, 15 g of bacteriological agar (Difco) was added.

### Growth conditions

Each mutant strain was grown on YPD at 30 °C with or without nourseothricin (100 μg/mL) or neomycin (200 μg/mL) as required for selection. After 2 d of culture, the cells were recovered and stored in glycerol stocks. For the experiment setup, strains were streaked on YPD plates and incubated for 2 d at 30 °C. A single colony was then used to inoculate 4 mL of YPD in a 13 mL round-bottom tube (Falcon) to generate a pre-inoculum. Cultures were incubated 2 d at 30 °C in a wheel rotor (70 rpm) to allow growth. Yeast cells were subsequently pelleted by centrifugation, washed twice with phosphate-buffered saline (PBS, pH 7), and centrifuged for 2 min at 2300 x *g*. Cell density was determined using a hematocytometer (Marienfeld, Lauda-Königshofen, Germany) and an Olympus BX40 microscope (Olympus, Hachioji, Tokyo) at 40X. This methodology was used every time prior to setting up the experiments.

For growth kinetic assays, 4 mL of the different fresh media was inoculated with 10^6^ yeast cells per mL. Growth was monitored by counting yeast cells daily for the time indicated.

### Thermotolerance assay, cell size, and capsule measurements

For thermotolerance assays, yeast cells were recovered from different liquid media after 4 d of growth, counted, washed, and then spotted in their respective solid media using serial dilutions (10^5^-10^1^ cells per spot). Plates were incubated at 30, 37, and 42 °C for 8 d. For capsule analysis, yeast cells grown in different media for 4 d at either 30 °C or 37 °C were suspended in India Ink for visualization. To induce capsule formation, yeast cells were grown in minimal media (MM) containing 15 mM glucose, 13 mM glycine, 10 mM MgSO_4_, 29.4 mM KH_2_PO_4_, and 0.3 µM thiamine. When indicated, alternative nitrogen and carbon sources were used at equimolar concentrations to the standard MM. A total of 4 x 10^6^ yeast cells were used for sample imaging, and pictures were acquired at 100X magnification.

### Cell wall and capsule fluorescent staining

Yeast cells grown in the different liquid media for 4 d at 30 °C or 37 °C were recovered and washed. A total of 25 x 10^6^ cells were incubated for 12 h at 4 °C in 100 μL of PBS containing 1% albumin and 100 μg/mL of 18B7 antibody [57], which binds the cryptococcal polysaccharide, reflecting indirect glucuronoxylomannan (GXM) content. Afterwards, cells were recovered, washed three times with PBS by centrifugation for 2 min at 2300 x *g*, and resuspended in 100 μL of PBS with 100 μg/mL of Alexa Fluor 488 (Thermo Fisher Scientific, Massachusetts, USA) conjugate. After an additional 12 h at 4 °C, cells were recovered, washed under the same conditions, and resuspended in 100 μL of PBS and 100 μg/mL of Uvitex 2B to stain chitin. Following a 2 h incubation at 4 °C, cells were recovered and washed under the same conditions, resuspended in 100 μL of 50 % glycerol in PBS. A total of 4 x 10^6^ yeast cells were used for sample visualization, and pictures were acquired at 100X magnification.

### Melanization, urease activity, and biofilm formation assays

The yeast cells, previously grown for 4 d on the different media, were recovered, washed, and used for downstream assays at either 30 °C or 37 °C. Melanization rate was assessed by spotting cells in minimal media (MM) containing 1 mM L-DOPA. For the urease activity assay, 1 x 10^8^ cells were inoculated in a round-bottom tube with 1 mL of urea liquid medium, containing 2 % urea (Sigma), 0.1 % glucose, 0.1 % peptone (Difco), 0.2% K_2_HPO_4_, 0.5 % NaCl (Sigma), and 0.0012 % phenol red (Sigma). Cultures were incubated with agitation, and aliquots were recovered at the indicated time points. Absorbance was measured at 560 nm using a SpectraMax iD5e plate reader.

For the biofilm formation assay, 2 x 10^5^ cells from YPD were transferred to a polystyrene 96-well plate (Corning) in 200 μL of PBS. After 1.5 h of adhesion, PBS was removed and replaced with 200 μL of the different fresh media. Plates were incubated for 72 h, washed with PBS, and air-dried for 45 min at room temperature. Biofilms were stained with 110 μL of 0.4 % violet crystal for 45 min, washed 4 times with PBS, and destained with 110 μL of 95 % ethanol for 45 min. The ethanol solution was transferred to a new plate, and its absorbance was measured at 595 nm on a SpectraMax iD5e.

### Macrophage-interaction assays

The J744 cell line or bone marrow-derived macrophages (BMDMs) were harvested and maintained according to previously reported protocols [44, 58]. For interaction assays [59], *C. neoformans* strains were grown in indicated media for either 1 or 4 d. Activated macrophages (5×10^5^) and *C. neoformans* yeast cells (15×10^5^) were co-inoculated into a 12-well plate containing 0.5 mL of Dulbecco’s Modified Eagle Medium (DMEM) for 2 h. The monoclonal 18B7 antibody was used as an opsonin at a final concentration of 10 μg/mL. For CFU determination, recovered cells were plated on Sabouraud plates and incubated for 48 h at 30 °C. Mitochondria were stained using 200 nM MitoTracker Red (Thermo Fisher Scientific). For peroxisome analysis, the AI100 strain was used, which expresses the DsRED protein with a peroxisome-targeting signal peptide [30]. Mitochondria and peroxisomes quantification was estimated by counting individual fluorescent dots per yeast cell, independent of size or fluorescence intensity.

### MitoTracker staining

The yeast cells, previously grown for 2 d on liquid YPD, were recovered, washed, and counted. Cells were inoculated at a density of 10^6^ cells/mL into 4 mL of minimal media supplemented with either 200 nM MitoTracker Red or MitoTracker Green (Thermo Fisher Scientific). The tubes were incubated for 2 d. A total of 4×10^6^ cells were used for visualization; images were acquired every 12 h for 2 d at 100X magnification. Mitochondrial quantification was estimated by counting individual fluorescent dots per yeast cell, independent of size or fluorescence intensity.

### Antioxidant treatment

Yeast cells previously grown for 2 d on liquid YPD were recovered, washed, and counted. Then, 10^6^ cells/mL were inoculated into 4 mL of MM supplemented with 1.5 mM of glutathione reduced (Sigma), ascorbic acid (Sigma), or H_2_O_2_ (Sigma).

Cultures were incubated at 30 °C for 2 d. Following antioxidant treatment, cells were used for capsule measurements and urease activity assay as described above. For the melanization assay, 4 mL of liquid MM + 1 mM of L-DOPA + antioxidants was inoculated with 10^6^ cells/ mL, and 0.5 mL aliquots were taken for imaging.

### Antifungal treatment

The yeast cells, previously grown for 2 d on liquid YPD, were recovered, washed, and counted. Then, 10^6^ cells/mL were inoculated into 4 mL of MM supplemented with 1.5 mM glutathione or ascorbic acid. Cultures were incubated at 30 °C with agitation for 2 d. Subsequently, cells were recovered, washed, and counted. A total of 10^6^ cells were transferred to a 96-well plate containing 200 μL of fresh MM supplemented with either 10 μg/mL of fluconazole (Sigma) or caspofungin (Sigma). The plate was incubated at 30 °C for 24 h, after which CFUs were determined on YPD agar.

### Mammalian tissue-derived nutrient’s agar

Fresh beef liver or heart (25 g) from the grocery store was finely sliced and resuspended with 30 mL of PBS without calcium and magnesium (Corning). For nutrient recovery, the liver solution was gently agitated upside down for 10 min, whereas the heart solution was vortexed for three cycles of 30 s. Homogenates were then centrifuged at 2300 x *g* for 10 min. The homogenates were spun down, and the process was repeated with fresh PBS. The resulting 50 mL of extract (nutrient solution) from the liver of the heart was diluted in 200 mL of PBS, filter-sterilized, and used to prepare 500 mL of agar (1.5%) plates. PBS-agar plates were used as a negative control, and MM-glucose agar served as a positive control.

### Galleria mellonella infection assays

*C*. *neoformans* was grown for 2 d in YPD, and then inoculated at 10^6^ cells/mL into 13 mL bottom-round tubes containing 4 mL of standard MM, MM with non-fermentable carbon sources, or MM supplemented with 1.5 mM antioxidants. Cultures were incubated with agitation at 30 °C for 4 d. Yeast cells were then recovered, washed, and counted. *G*. *mellonella* larvae were obtained from Vanderhorst Wholesale (Ohio, USA). Final instar *G*. *mellonella* larvae weighing 175-225 mg were selected. Groups of 10 larvae were injected with either 10 μL of PBS or 10 μL of 10^5^ *C*. *neoformans* yeast cells in PBS per larva. *G*. *mellonella* larvae and pupae were kept at 30°C and monitored daily for survival for 14 d. Survival was assessed by movement in response to a pipette stimulus.

### Image analysis

Images were acquired using QCapture software with an Olympus AX70 microscope (Olympus, Tokyo, Japan) at a 100X objective. Cell size, capsule thickness, and fluorescence intensity were measured using Fiji software [60].

### Statistical analysis

Graph Prism 10.6.1 software was used to create the plots and statistical analysis. Data were analyzed using one-way ANOVA with Tukey’s post-hoc test, ****P<0.0001. For survival data analysis, the Mantel-Cox test was used, * P<0.05, ** P<0.01.

## Author contributions

JA-PM: Conceptualization, Methodology, Validation, Formal analysis, Investigation, Writing - Original Draft, Writing - Review & Editing, Visualization; EC: Methodology, Writing - Review & Editing; AC: Conceptualization, Resources, Writing - Review & Editing, Supervision, Project administration, Funding acquisition.

## Acknowledgments

We thank Kronstad and Heitman labs for providing the mutant strains used in this study. We thank Piotr Stempinski for technical support during biofilm assays. We thank Seth Greengo, Quigly Dragotakes, Samuel Rodrigues dos Santos, and Sebastien Ortiz for technical support during the macrophage experiments. We thank Daniel Smith for technical support during *G. mellonella* assays. This work was supported by NIH grants AI052733–16, AI152078–01, and HL059842–19.

## Data availability

All data generated in this study are provided in the manuscript.

## Supporting information

This article contains supporting information.

