## Supplemental material for "Metabolic plasticity and virulence of *Cryptococcus neoformans* are regulated by mitochondrial homeostasis"

**Table1. Strains used in this study**

| Strain | Genotype | Reference |
|---|---|---|
| *C. neoformans* var. *grubii* H99 | WT strain (serotype A, MATα). | [#]Perfect *et al*., 1993. |
| *hap3*Δ | *MATα hap3Δ::NAT* | Jung *et al*., 2010 [31]. |
| *hap5*Δ | *MATα hap5Δ::NAT* | Jung *et al*., 2010 [31]. |
| *hapX*Δ | *MATα hapXΔ::NAT* | Jung *et al*., 2010 [31]. |
| *hap3*Δ::*hap3* | *MATα hap3Δ::NAT, HAP3-NEO* | Jung *et al*., 2010 [31]. |
| *hap5*Δ::*hap5* | *MATα hap5Δ::NAT, HAP5-NEO* | Jung *et al*., 2010 [31]. |
| *hapX*Δ::*hapX* | *MATα hapXΔ::NAT, HAPX-NEO* | Jung *et al*., 2010 [31]. |
| *pka1*Δ-1 (YSB188) | *MAT*α *pka1*Δ::*NAT-STM#191* | Bahn *et al*., 2004 [31]. |
| *pka1*Δ-2 (YSB189) | *MAT*α *pka1*Δ::*NAT-STM#192* | Bahn *et al*., 2004 [31]. |
| *pka2*Δ-1 (YSB194) | *MAT*α *pka2*Δ::*NAT-STM#205* | Bahn *et al*., 2004 [31]. |
| *pka2*Δ-2 (YSB195) | *MAT*α *pka2*Δ::*NAT-STM#206* | Bahn *et al*., 2004 [31]. |
| *C. neoformans* var. *grubii* KN99 | WT strain (serotype A, MATα). | |
| AI100 (*C. neoformans* KN99) | *$PH_3$-DsRED-SKL-NEO MATα* | Idnurm *et al*., 2007 [30]. |

Kronstad Lab kindly provided the *hap* mutant strains.

Heitman Lab kindly provided the *pka* mutant and AI100 strains.

### Perfect JR, Ketabchi N, Cox GM, Ingram CW, Beiser CL (1993) Karyotyping of *Cryptococcus neoformans* as an epidemiological tool. *J Clin Microbiol* 31: 3305-3309.

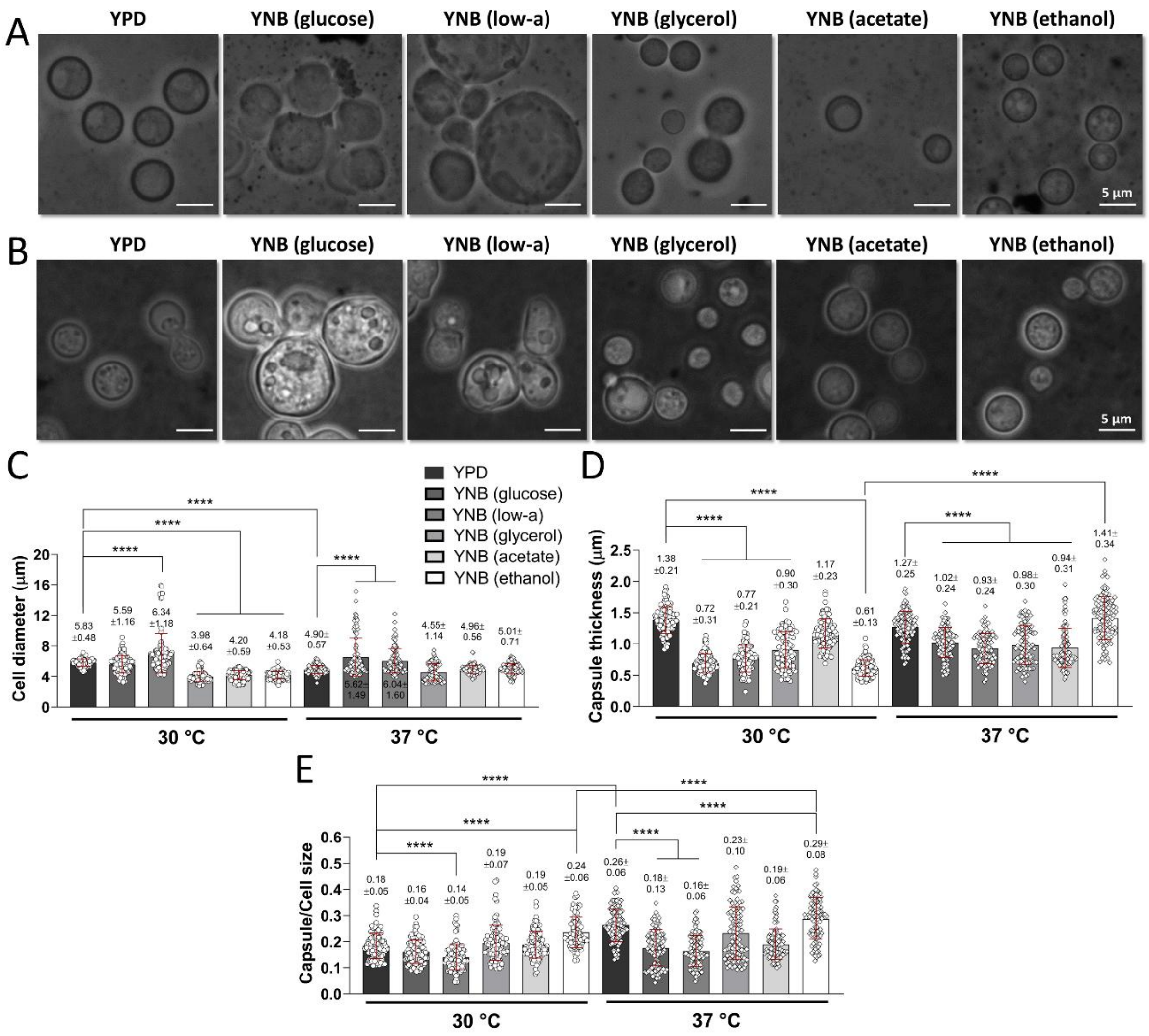


**Figure S1. The carbon source impacts cell morphology and capsule thickness of *C. neoformans*. (A)** Representative microphotographs of India Ink staining of H99 strain grown on YNB with non-fermentable carbon sources at 30 °C (day 4). **(B)** Representative microphotographs of India Ink staining of H99 strain grown on YNB with non-fermentable carbon sources at 37 °C (day 4). **(C)** Quantification of cell diameter (cell size) from panels A and B. **(D)** Quantification of the capsule thickness from panels A and B. **(E)** Quantification of the ratio between capsule and cell size from panels A and B. Plots show individual measurements of three biological replicates with technical duplicates, n=100, mean ± SD. One-way ANOVA with Tukey test, **** p<0.0001.

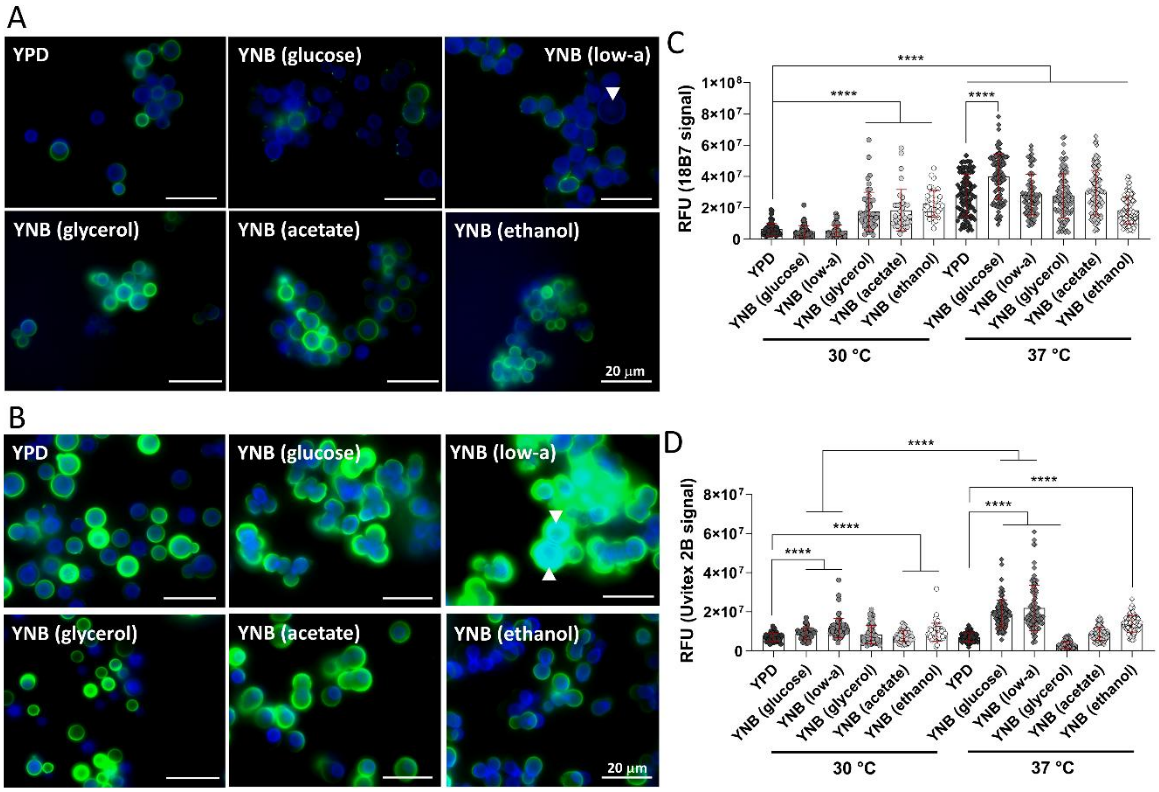


**Figure S2. Using a non-fermentable carbon source increases the levels of glucuronoxylomannan (GXM) and chitin of *C. neoformans*. (A)** Representative microphotographs of fluorescent staining of H99 strain grown on YNB with non-fermentable carbon sources at 30 °C (day 4). **(B)** Representative microphotographs of fluorescent staining of H99 strain grown on YNB with non-fermentable carbon sources at 37 °C (day 4). The green signal denotes GXM, while the blue signal denotes chitin. **(C)** Quantification of GXM fluorescence intensity from panels A and B. **(D)** Quantification of chitin fluorescence intensity from panels A and B. Plots show individual measurements of three biological replicates with technical duplicates, n=100, mean ± SD. One-way ANOVA with Tukey test, **** p<0.0001.

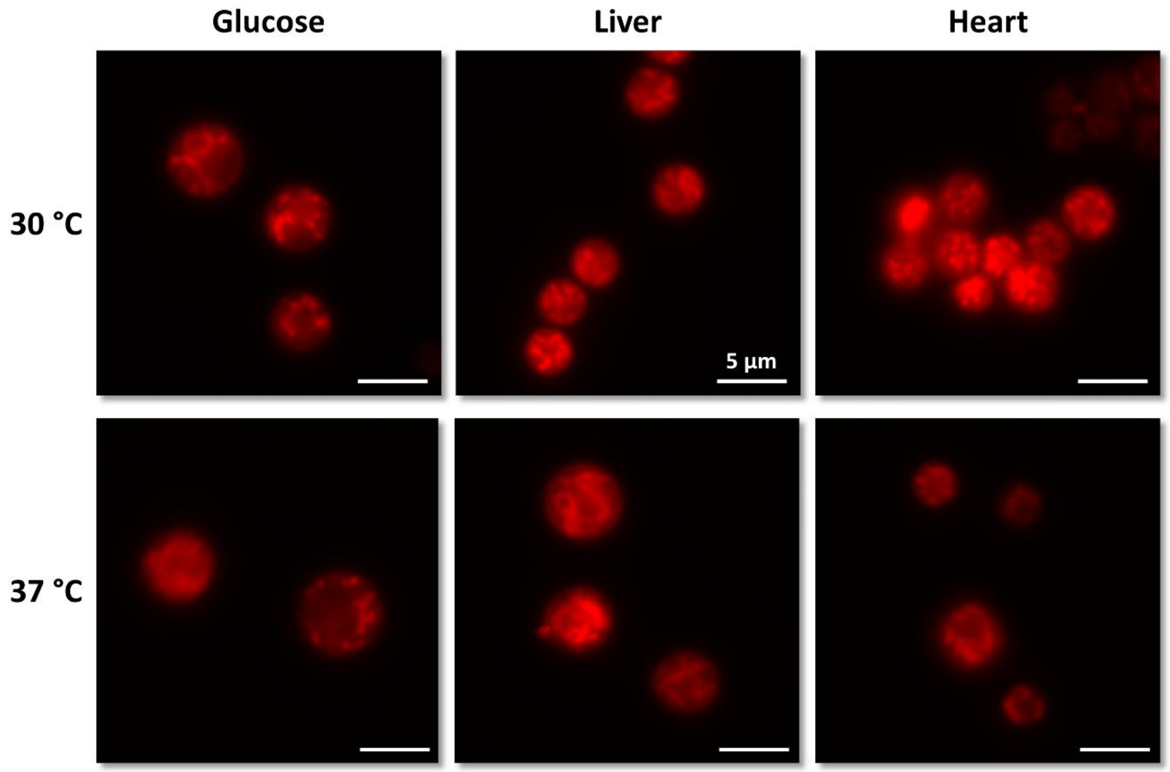


**Figure S3. Mitochondrial activity is enhanced under a host-tissue-mimicking condition.** Representative microphotographs of H99 strain stained with MitoTracker Red after 5 d of growth on the liver or heart nutrients agar at the indicated temperatures, 100X magnification.

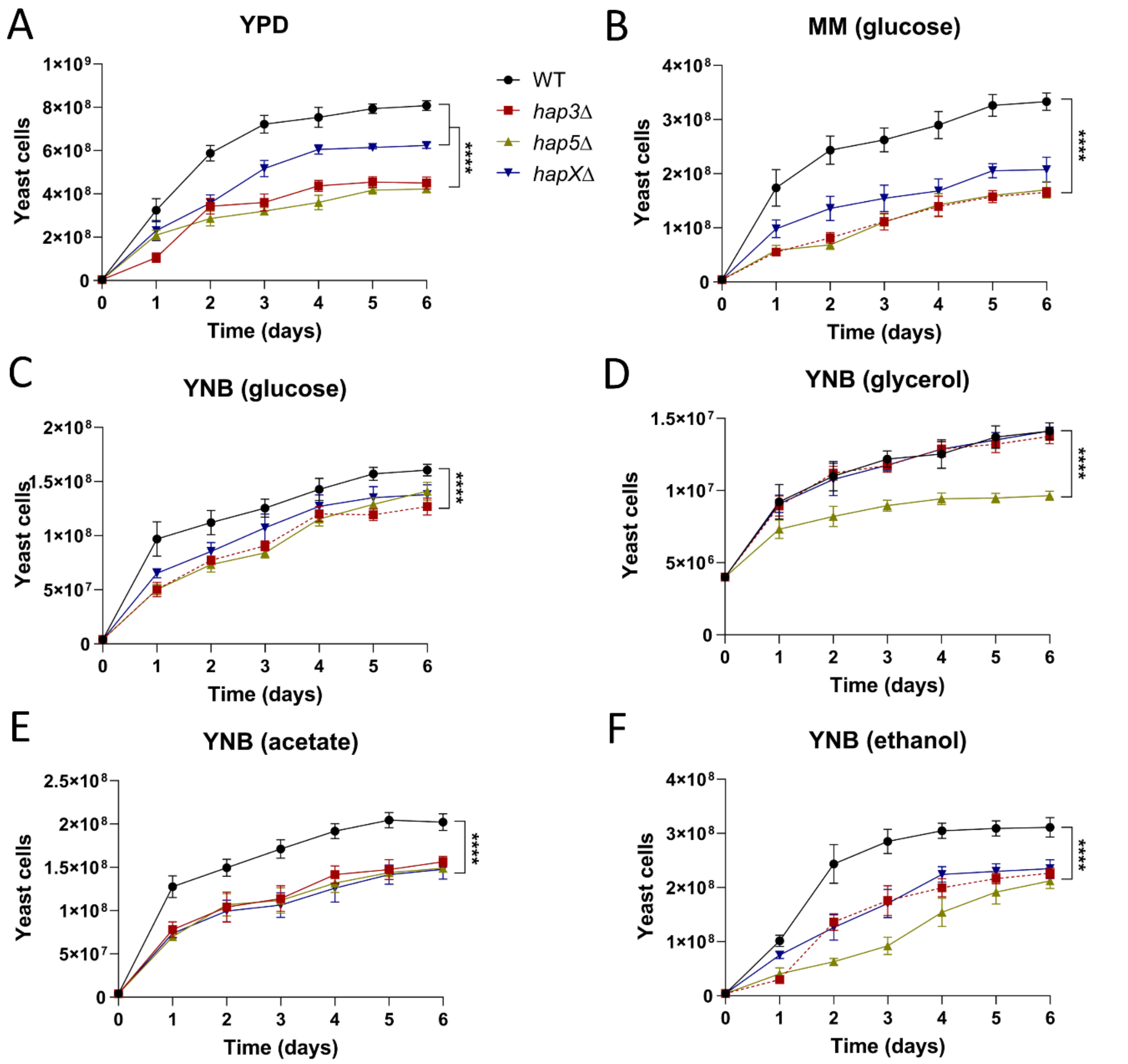


**Figure S4. The deletion of Hap complex genes impairs growth. (A)** Liquid growth kinetics of H99 and *hap*Δ mutant strains in rich medium YPD. **(B)** Liquid growth kinetics of H99 and *hap*Δ mutant strains in minimal medium MM-glucose. **(C)** Liquid growth kinetics of H99 and *hap*Δ mutant strains in minimal medium YNB-glucose. **(D)** Liquid growth kinetics of H99 and *hap*Δ mutant strains in minimal medium YNB-glycerol. **(E)** Liquid growth kinetics of H99 and *hap*Δ mutant strains in minimal medium YNB-acetate. **(F)** Liquid growth kinetics of H99 and *hap*Δ mutant strains in minimal medium YNB-ethanol. Plots show the average of three biological replicates with technical duplicates, n=6, mean ± SD. One-way ANOVA with Tukey test, **** p<0.0001.

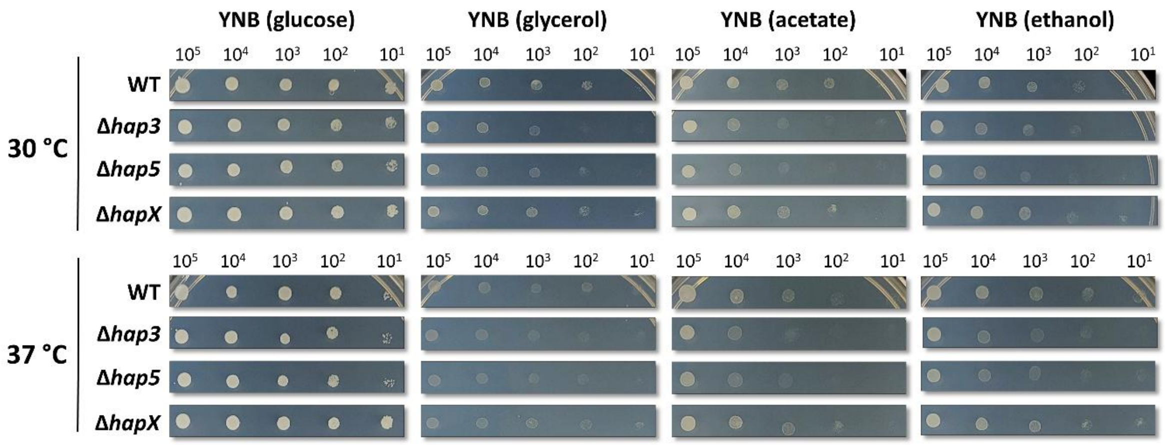


**Figure S5. The deletion of Hap complex genes decreased thermotolerance.** Representative photos at day 8 of H99 and *hap*Δ mutant strains growth on solid media with non-fermentable carbon sources at the indicated temperatures.

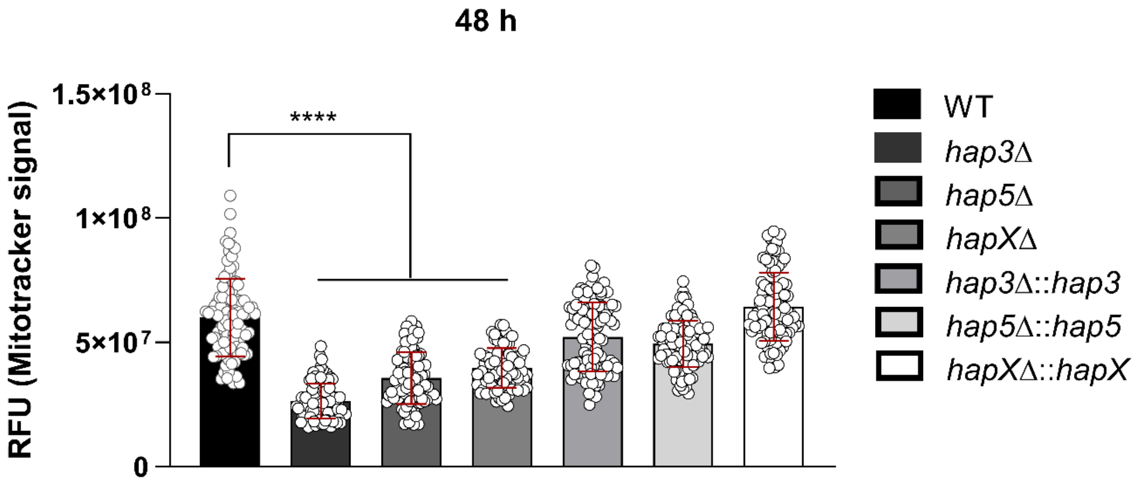


**Figure S6. The deletion of Hap complex genes reduced mitochondrial activity.** Quantification of mitochondrial activity (fluorescent intensity) after 48 h of growth on MM-glucose of H99 and *hap*Δ mutant strains stained with MitoTracker Red. Plot shows the average of three biological replicates with technical duplicates, n=100, mean ± SD. One-way ANOVA with Tukey test, **** p<0.0001.

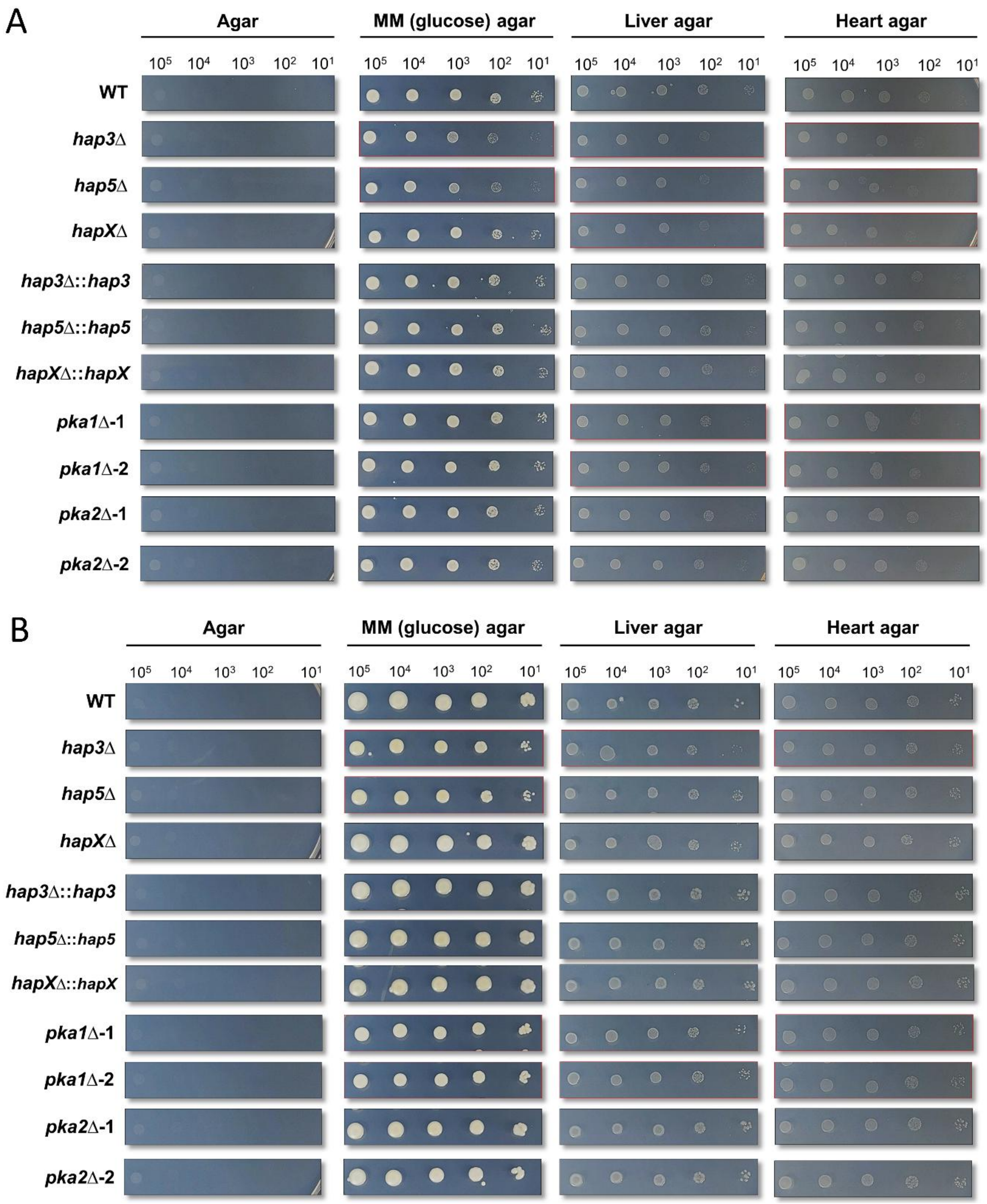


**Figure S7. The deletion of *hap* complex and *pka1* genes impairs growth in a tissue-mimicking condition. (A)** Representative photos at day 2 of H99, *hap*Δ, *hap*-complemented, and *pka*Δ mutant strains grown on solid media with nutrients from liver or heart at 30 °C. **(B)** Representative photos at day 4 of H99, *hap*Δ, *hap*-complemented, and *pka*Δ mutant strains grown on solid media with nutrients from liver or heart at 37 °C.

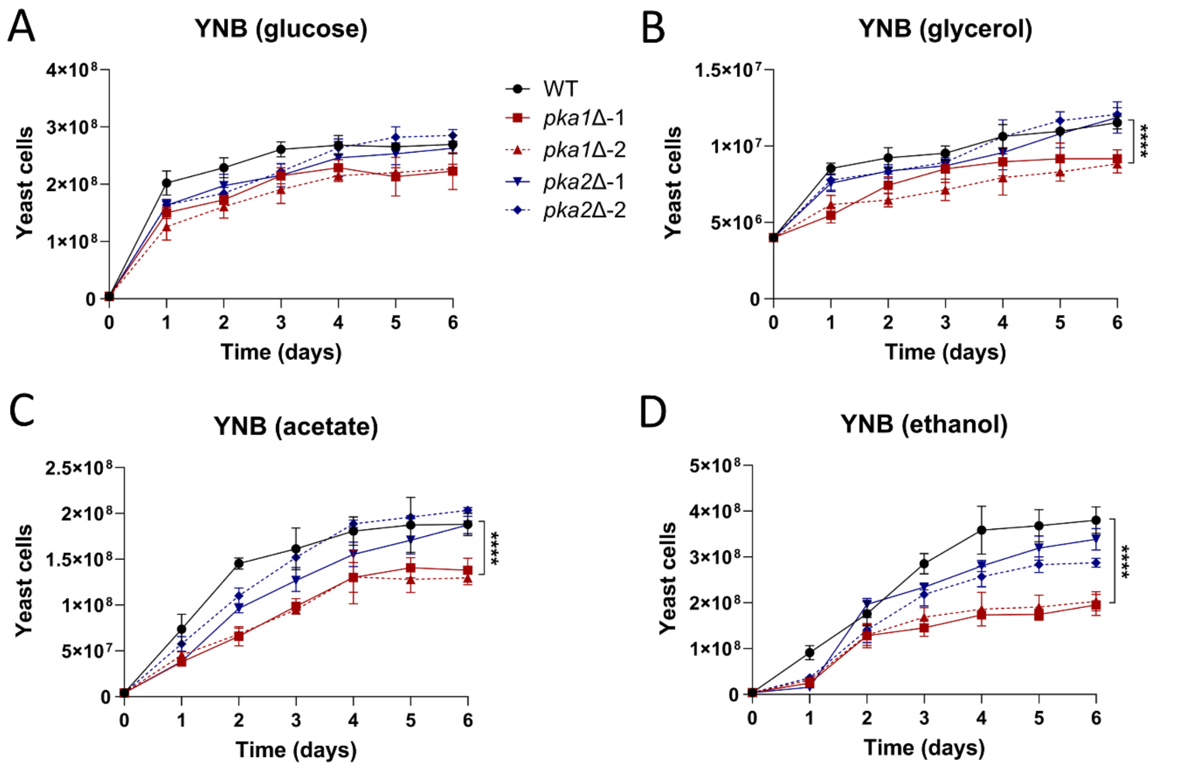


**Figure S8. Deletion of the *pka1* gene impairs growth on non-fermentable carbon sources. (A)** Liquid growth kinetics of H99 and *pka*Δ mutant strains in minimal medium YNB-glucose. **(B)** Liquid growth kinetics of H99 and *pka*Δ mutant strains in minimal medium YNB-glycerol. **(C)** Liquid growth kinetics of H99 and *pka*Δ mutant strains in minimal medium YNB-acetate. **(D)** Liquid growth kinetics of H99 and *pka*Δ mutant strains in minimal medium YNB-ethanol. Plots show the average of three biological replicates with technical duplicates, n=6, mean ± SD. One-way ANOVA with Tukey test, **** $p<0.0001$.

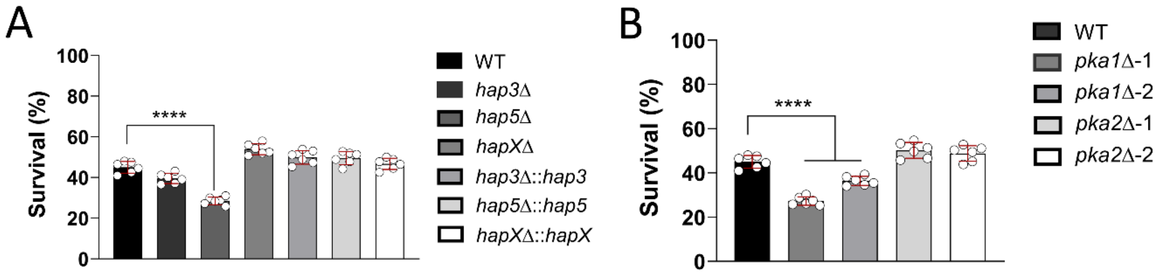


**Figure S9. Hap5 and Pka1 are key components for survival and resistance to macrophage phagocytosis. (A)** Quantification of survival of yeast cells of H99 and *hap*Δ mutant strains grown on MM-glucose and challenged with J774 macrophages. **(B)** Quantification of survival of yeast cells of H99 and *pka*Δ mutant strains grown on MM-glucose and challenged with J774 macrophages. Plots show the average of three biological replicates with technical duplicates, n=6, mean ± SD. One-way ANOVA with Tukey test, **** p<0.0001.
